# METTL3 inhibition-activated cGAS/STING axis enhances immunotherapy and PARP inhibitor sensitivity of lung carcinoma

**DOI:** 10.1101/2024.09.03.610931

**Authors:** Jiawang Zhou, Jiaxin He, Yunqing Lu, Cheng Yi, Xing Chang, Lijun Tao, Ke Zhong, Haisheng Zhang, Jiexin Li, Zhuojia Chen, Hongsheng Wang

## Abstract

The cGAS/STING-mediated type I interferon response can augment antitumor activity, while the regulatory factors within this innate immune response remain elusive. Herein we found that the RNA m^6^A methyltransferase METTL3 was upregulated in lung carcinoma tissues. Elevated METTL3 level was correlated with diminished CD8^+^ T cell infiltration and cancer progression in lung carcinoma patients. METTL3 deficiency exacerbated nuclear DNA leakage into the cytoplasm, activating the cGAS pathway and thereby enhancing anti-tumor immunity. Mechanistically, METTL3 deficiency reduced the homologous recombination repair efficacy via downregulation of MSH5, a mutS family protein involved in DNA mismatch repair, leading to increased cytosolic DNA levels. m^6^A methylation of A2521 of *MSH5* stabilized its mRNA via binding with IGF2BP2. On the other hand, m^6^A methylation of A1545 at the CDS of cGAS decreased mRNA stability and regulates its protein expression. Functionally, knockdown of METTL3 sensitized lung carcinoma cells to the PARP inhibitors. *In vivo* and clinical data confirmed the positive roles of METTL3 inhibition-activated cGAS/STING axis in tumor growth and lung adenocarcinoma progression. Collectively, METTL3 inhibition activates the cGAS/STING-mediated anti-tumor immunity via induction of cytosolic DNA and cGAS expression, which in turn regulate PARP inhibitor response and cancer progression in lung carcinoma.

## Introduction

Lung carcinoma, a prevalent and lethal malignancy within the respiratory system, ranks at the forefront of cancer incidence and mortality rates worldwide, asserting a significant public health concern^[1]^. In recent years, immunotherapy has emerged as a groundbreaking modality for lung carcinoma treatment, marking an epochal advancement in the annals of oncology^[2]^. Nonetheless, the heterogeneous pathological types, drug resistance, and recurrence of lung carcinoma present formidable challenges to effective treatment. Chief among these are immunosuppression and treatment resistance challenges that immunotherapeutic approaches grapple with. This necessitates the incorporation of immunotherapy into a multimodal treatment paradigm, capitalizing on synergistic effects to enhance clinical efficacy and mount a more targeted offensive against tumors, which represents the current cutting-edge in cancer therapeutic research^[3]^. Hence, delving into innovative strategies that synergistically augment immunotherapy for lung carcinoma has become a critical and urgent priority for scientific inquiry.

*N^6^*-methyladenosine (m^6^A) is the most prevalent internal modification on eukaryotic RNAs^[4]^. Increasing evidence supports that m^6^A, along with its regulatory enzymes and reader proteins play critical roles in post-transcriptional gene regulation, and their dysregulation has been associated with various diseases, especially cancer^[5, 6]^. Methyltransferase-like 3 (METTL3), one of the most studied and well-known m^6^A writer, serves as a key component of methyltransferases complex (MTC) and is crucial in the regulation of m^6^A modification^[7]^, METTL3 has been implicated in both oncogenic or tumour-suppressive roles in multiple cancers^[8–10]^. Inhibition of METTL3 has been shown to reprogram tumor microenvironment (TME) and enhance the effectiveness of anti-PD-1 treatment^[11]^. A deeper understanding of the role and the underlying molecular mechanisms of METTL3 in anti-cancer immunity will enhance the development of effective targeted therapeutics against METTL3.

DNA damage leads to the accumulation of cytosolic DNA fragments within cells^[12]^. The cytosolic accumulated DNA may induce the production of type I interferon (IFN) by activation of cyclic GMP-AMP synthase (cGAS) / Stimulator of interferon genes (STING) signaling pathway. This activation occurs through the binding of cytosolic DNA to cGAS, which then catalyzes the production of the second messenger cGAMP^[13]^. Subsequently, cGAMP interacts with and activates STING, leading to the the recruitment and activation of TANK-binding kinase 1 (TBK1), triggers the interferon regulator factor 3 (IRF3) and NF-κB signaling cascades and cytokines production^[14]^. The reactivation of cGAS/STING signaling pathway has been shown to sensitize cancer cells to chemotherapy. Combination of STING agonist with carboplatin can reduce tumor burden and extend survival in mouse model^[15]^.

The post-transcriptional regulation of cGAS/STING signaling pathway has been extensively studied^[16]^. It has been reported that TRIM14 recruits USP14 to cleave the lysine 48 (K48)-linked ubiquitin chains of cGAS at K414, thereby inhibiting p62-mediated autophagic degradation of cGAS to enhance the activation of type I interferon signaling^[17]^. While USP35 can directly deubiquitinate and inactivate STING by phosphorylation of STING at Ser366^[18]^. However, there is limited evidence connecting cGAS/STING signaling pathway and epigenetic regulatory factors. Whether m^6^A modification could directly regulate cGAS/STING pathways and affect anti-tumor immunity remain unknown. In the present study, we investigated the potential effects of m^6^A on activating cGAS-mediated anti-tumor immunity of lung carcinoma cells. Our data revealed that METTL3 inhibition activates the cGAS-mediated anti-tumor immunity via induction of cytosolic DNA and cGAS expression, which in turn regulate PARP inhibitor response and cancer progression in lung carcinoma.

## Results

### 1. METTL3 expression is correlated with immune cell infiltration and cancer progression

To elucidate the potential pathological roles of METTL3 in the anti-tumor immunity of lung adenocarcinoma, we first investigated the correlation between METTL3 expression and immune cell infiltration via the TIMER2.0 web server (http://timer.comp-genomics.org/). Results revealed that METTL3 expression was negatively correlated with the infiltration of diverse anti-tumor immune cells, including CD8^+^ T cells, CD4^+^ T cells (Figure 1 A), M1 macrophages, monocytes, and neutrophils (Figure S1 A). Conversely, the m^6^A demethylase, FTO displayed positive correlations with the infiltration of these anti-tumor immune cells (Figure 1 B). In addition, METTL3 expression was positively correlated with the infiltration of pro-tumor immune cells in lung adenocarcinoma, including M2 macrophages and regulatory T cells (Tregs) (Figure 1 C). Notably, similar correlations were observed in several other types of cancers (Figure S1 B), suggesting a broader implication of METTL3 in modulating immune cell infiltration in tumor microenvironments.

**Figure 1.**
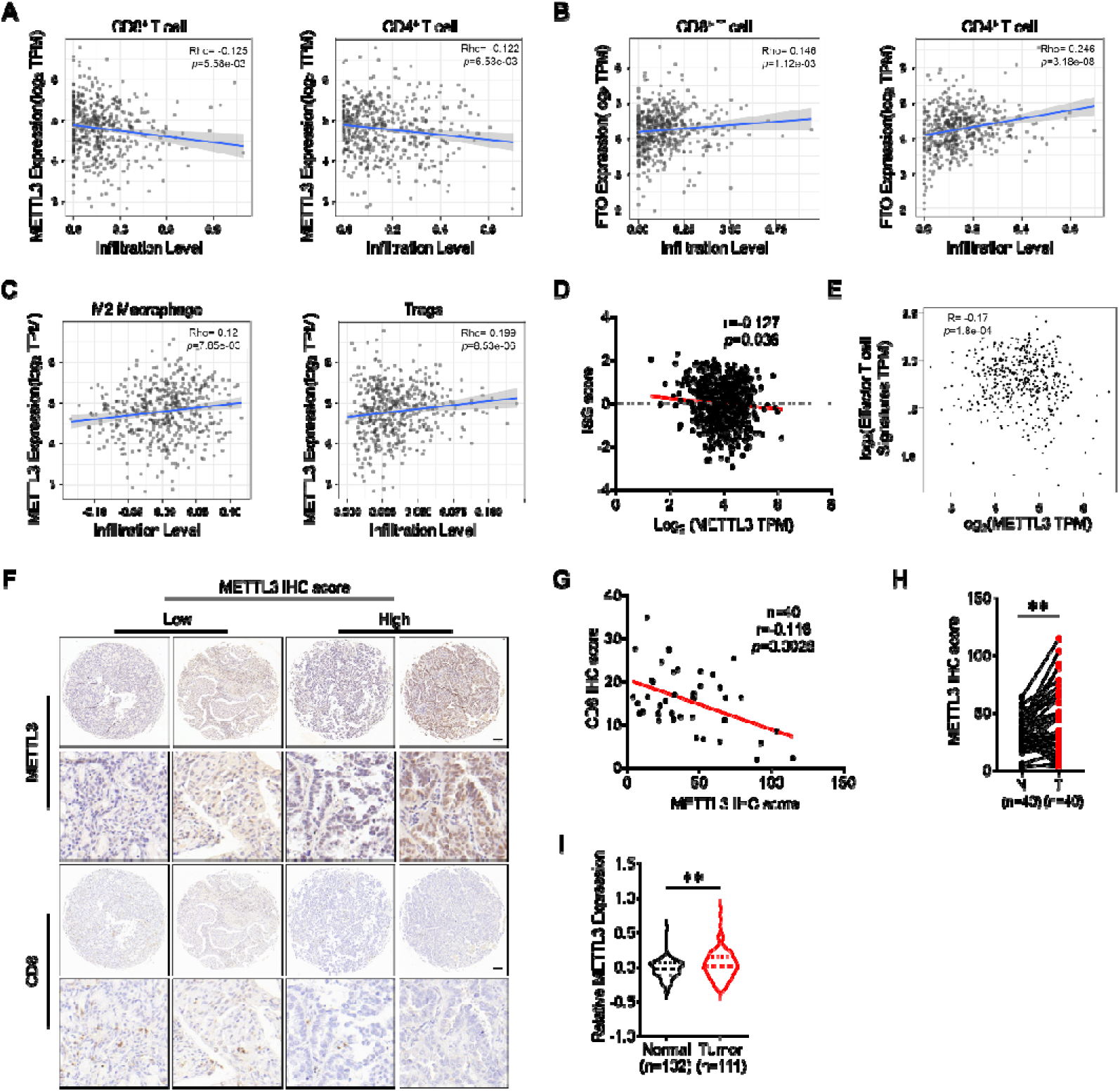
METTL3 expression is correlated with immune cell infiltration and cancer progression. (A) The correlation between the expression of METTL3 and infiltrating level of CD8^+^ T cells or CD4^+^ T cells in lung adenocarcinoma based on TIMER platform; (B) The correlation between the expression of FTO and infiltrating level of CD8^+^ T cells or CD4^+^ T cells in lung adenocarcinoma based on TIMER platform; (C) The correlation between the expression of METTL3 and infiltrating level of M2 macrophages or Tregs in lung adenocarcinoma based on TIMER platform; (D) The correlation between the expression of METTL3 and ISG scores in lung adenocarcinoma based on TCGA database; (E) The correlation between the expression of METTL3 and effector T cell markers (including CX3CR1, FGFBP2, FCGR3A) in lung adenocarcinoma based on TCGA database; (F) Representative images of CD8^+^ T cells from high or low IHC scores of METTL3 in lung adenocarcinoma tissues, scale bar = 100 μm; (G) The correlation between the IHC scores of METTL3 and the IHC scores of CD8 in lung adenocarcinoma tissues; (H) The IHC scores of METTL3 in LUAD tumor tissues and adjacent normal mucosa tissues; (I) The expression of METTL3 in LUAD tumor tissues and adjacent normal mucosa tissues from CPTAC database; Data are presented as mean ± SD from three independent experiments. \**p*<0.05, \*\**p*<0.01, \*\*\**p*<0.001, ns, no significant, by Student’s *t* test between two groups and by one-way ANOVA followed by Bonferroni test for multiple comparison.

Utilizing a previously established method for calculating the ISG score based on the expression profile of a 38-gene signature^[19]^, we observed that high expression of METTL3 in lung adenocarcinoma was associated with a lower ISG score (Figure 1 D). Additionally, the expression of effector T cell markers (including CX3CR1, FGFBP2, FCGR3A)^[20]^ were negatively correlated with METTL3 expression (Figure 1 E), but positive correlated with FTO expression (Figure S1 C) in lung adenocarcinoma. Furthermore, immunohistochemical (IHC) staining results revealed a significant reduction in intratumoral CD8^+^ cell infiltration in lung adenocarcinoma specimens with high METTL3 expression (Figure 1 F and S1 D), the protein expression of METTL3 was negatively correlated with CD8^+^ cell infiltration (Figure 1 G). These findings indicate that METTL3 may contribute to immune regulation within lung adenocarcinoma.

Beyond lung adenocarcinoma, METTL3 expression was also upregulated in several other cancer types, including colon adenocarcinoma, esophageal carcinoma, and prostate adenocarcinoma, compared to their corresponding normal adjacent tissues (Figure S1 E). Consistently, the protein expression of METTL3 was significantly increased in lung adenocarcinoma patients from immunohistochemical staining results (Figure 1 H) and the CPTAC database (Figure 1 I). These findings indicate an inverse relationship between METTL3 expression and the infiltration of anti-tumor immune cells, as well as cancer progression in lung adenocarcinoma.

### 2. Knockdown of METTL3 promotes the innate immune response and activates the cGAS-STING pathway

To delve into the mechanism underlying how METTL3 regulates immune cell infiltration, we established sh-control and sh-METTL3 A549 and H1299 cell lines with lentiviruses (Figures S2 A). Given that cytoplasmic DNA is a primary activator of the innate immune response^[21]^, we proceed to examine DNA breaks and cytosolic DNA accumulation. To measure cytosolic DNA, we utilized PicoGreen dye, a widely used fluorescent stain that specifically binds to double-stranded DNA (dsDNA). Our observations revealed that under untreated conditions, the percentage of sh-METTL3 A549 cells with accumulated cytosolic DNA was significantly higher than that of sh-control A549 cells (Figures 2 A). This percentage increased substantially when cells were treated with Zeocin, a radio-mimetic chemical that predominantly induces DSBs^[22]^ (Figures 2 B) or exposed to 10 Gy of ionizing radiation (IR) (Figures S2 B). To confirm the essential roles of m^6^A in DNA breaks and cytosolic DNA accumulation, A549 cells were transfected with wild-type (WT) METTL3 and catalytically inactive METTL3 mutant DA (D395A)^[23]^ (Figures S2 C). Results revealed that the overexpression of METTL3, but not the METTL3 DA mutant, reversed the cytosolic DNA accumulation in A549 cells (Figures 2 C). These findings suggest that METTL3 deficiency leads to the release of DNA from the nucleus to the cytoplasm via an m^6^A-dependent manner.

**Figure 2.**
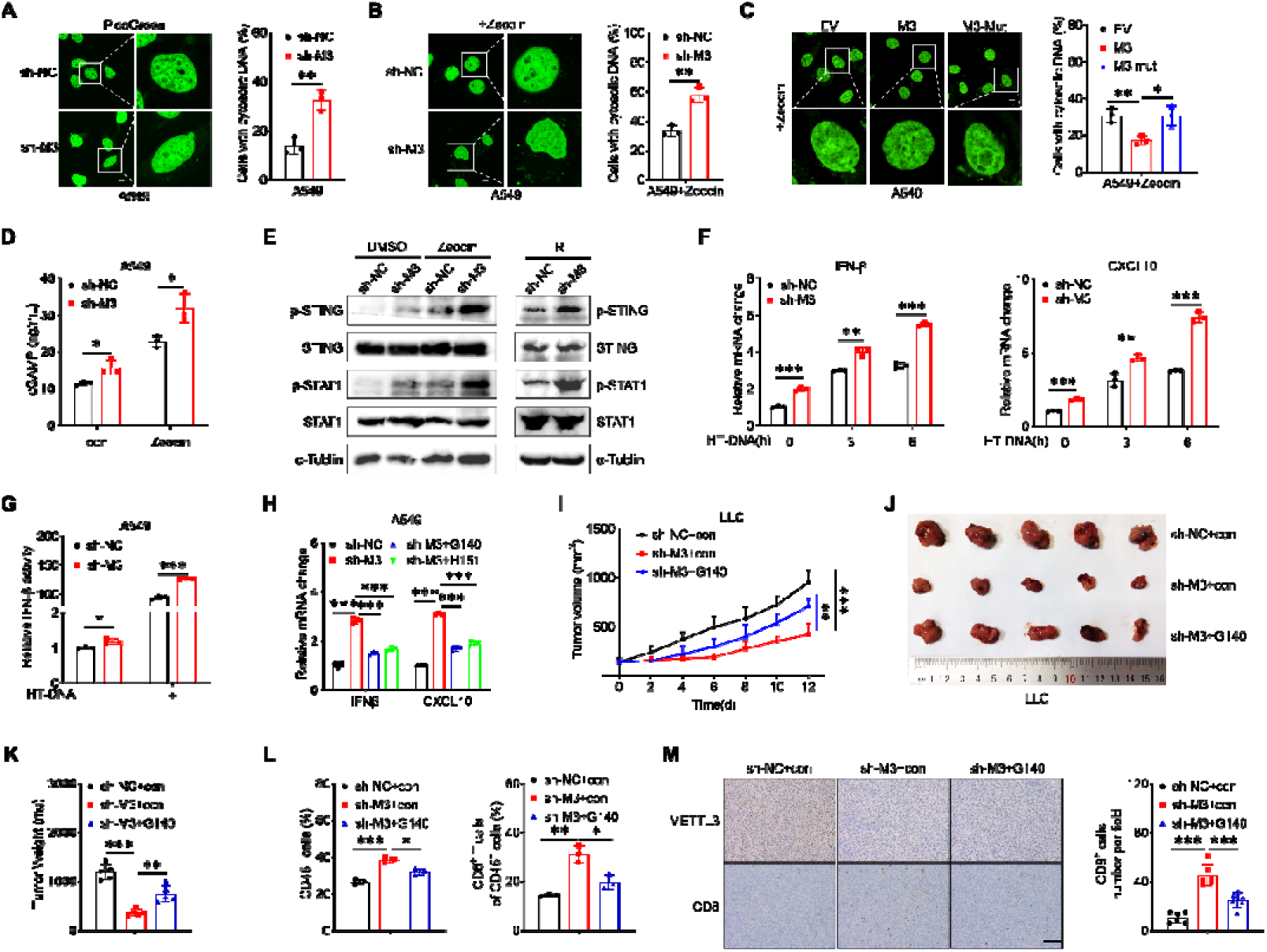
Knockdown of METTL3 promotes the innate immune response and activates the cGAS-STING pathway. (A) Representative confocal images of PicoGreen stain in sh-control and sh-METTL3 A549 cells (left) and the percentages of cells displaying cytosolic DNA were measured (right), scale bar = 10 μm; (B) Representative confocal images of PicoGreen stain in sh-control and sh-METTL3 A549 cells treated with Zeocin (left) and the percentages of cells displaying cytosolic DNA were measured (right), scale bar = 10 μm; (C) Representative confocal images of PicoGreen stain in A549 cells with Zeocin treatment transfected with vector control, METTL3 WT plasmid, METTL3 DA mutant plasmid for 24 h (left) and the percentages of cells displaying cytosolic DNA were measured (right), scale bar = 10 μm; (D) Intracellular cGAMP levels in sh-control and sh-METTL3 A549 cells with and without Zeocin treatment; (E) The protein expression of phosphorylation of STING (p-STING) and STAT1 (p-STAT1) in sh-control and sh-METTL3 A549 cells with or without Zeocin, IR treatment; (F) The mRNA levels of IFN-β, CXCL0 mRNA in sh-control and sh-METTL3 A549 cells treated with HT-DNA for 4 h; (G) The IFN-β promoter activities in sh-control and sh-METTL3 A549 cells treated with HT-DNA for 4 h; (H) The mRNA levels of IFN-β, CXCL0 mRNA in sh-control and sh-METTL3 A549 cells treated with G140 or H151 for 4 h; (I) The tumor growth curves of syngeneic tumor models using sh-control, sh-*METTL3* LLC cells with or without G140 treatment; (J) The tumor volume of syngeneic tumor models using sh-control, sh-*METTL3* LLC cells with or without G140 treatment at the end of the experiment; (K) The tumor weight of syngeneic tumor models using sh-control, sh-*METTL3* LLC cells with or without G140 treatment; (L) The percentages of CD45^+^ cell and the percentages of CD8^+^ T cell in CD45^+^ cell in the tumor tissues taken from mice with sh-control, sh-*METTL3* LLC syngeneic tumor with or without G140 treatment; (M) IHC (METTL3 and CD8)-stained paraffin-embedded sections obtained from sh-control, sh-*METTL3* LLC syngeneic tumor with or without G140 treatment, the scale bar is 100 μm. Data are presented as mean ± SD from three independent experiments. \**p*<0.05, \*\**p*<0.01, \*\*\**p*<0.001, ns, no significant, by Student’s *t* test between two groups and by one-way ANOVA followed by Bonferroni test for multiple comparison.

To determine whether the increased cytosolic DNA in sh-*METTL3* cells leads to cGAS activation, we measured the production of cyclic guanosine monophosphate (cGAMP) in sh-control and sh-*METTL3* lung adenocarcinoma cells. The results indicated that sh-METTL3 lung adenocarcinoma cells, regardless of Zeocin treatment, had significantly higher levels of cGAMP compared to sh-control cells, and Zeocin treatment further stimulated cGAMP production (Figure 2 D and S2 D). Consistent with the fact that cGAMP activates STING and its downstream factors, the increase in cGMAP in sh-*METTL3* A549 cells is associated with increased phosphorylation of STING and STAT1 (Figures 2 E). The interferon pathway is another downstream pathway of cGAS activation. We found that HT-DNA-triggered expression of IFN-β, as well as CXCL10, were significantly enhanced in sh-*METTL3* A549 cells (Figures 2 F). Furthermore, the luciferase reporter assay using pGL3-Basic-IFN-β-luc showed that the IFN-β promoter activity were significantly enhanced in sh-*METTL3* A549 cells (Figure 2 G). Consistent results were obtained by treatment of STM2457, a highly potent and selective inhibitor of METTL3/14 ^[24]^ (Figures S2 E and F). These findings indicate knockdown of METTL3 activates the cGAS-STING pathway.

The innate immune signaling triggered by *METTL3* deficiency depends on cGAS and STING, as METTL3 deficiency-activated interferon genes were abolished by either cGAS inhibitor G140^[25]^ or STING inhibitor H151^[26]^ treatment (Figures 2 H). To corroborate these findings *in vivo*, sh-control, sh-*METTL3* LLC cells were used to establish syngeneic tumor models. Results showed that knockdown of METTL3 significantly inhibited the growth of LLC xenografts, while the G140 treatment can rescue this effect (Figure 2 I). At the end of the experiment, tumor volume (Figure 2 J) and weight (Figure 2 K) in the sh-*METTL3* group were significantly lower than those measured in the sh-control group, and these reductions were reversed by G140 treatment. No significant change of body weight was observed among all groups (Figure S2 G). Further, flow cytometry analysis demonstrated that knockdown of METTL3 significantly increase the populations of CD45^+^ cell and the CD8^+^ T cell in CD45^+^ cell, while the G140 treatment can rescue this effect (Figure 2 L and S2 H). IHC analysis showed that sh-*METTL3* LLC tumor tissue exhibited high expression levels of CD8 compared to control tumor tissue (Figure 2 M). Collectively, these data indicate that METTL3 deficiency exacerbates the nuclear DNA leakage into the cytoplasm, thereby triggering anti-tumour immunity by activating the cGAS pathway.

### 3. METTL3 regulates cytosolic DNA levels by modulating homologous recombination repair (HR) efficacy

Considering that the accumulation of cytoplasmic DNA can be a consequence of nuclear DNA damage, we next assessed the phosphorylation of histone H2AX (γH2AX), an indirect marker of DNA double-strand breaks (DSBs) in lung adenocarcinoma cells. Western blot analysis showed that knockdown of METTL3 significantly elevated the γH2AX level in A549 and H1299 cells (Figure 3 A). Besides, γH2AX levels persisted in sh-*METTL3* A549 cells can be induced more strongly by Zeocin treatment (Figures 3 B). Additionally, the confocal analysis confirmed the same conclusion (Figures 3 C). These results suggest that sh-*METTL3* cells have delayed repair and/or persistent generation of DSBs.

**Figure 3.**
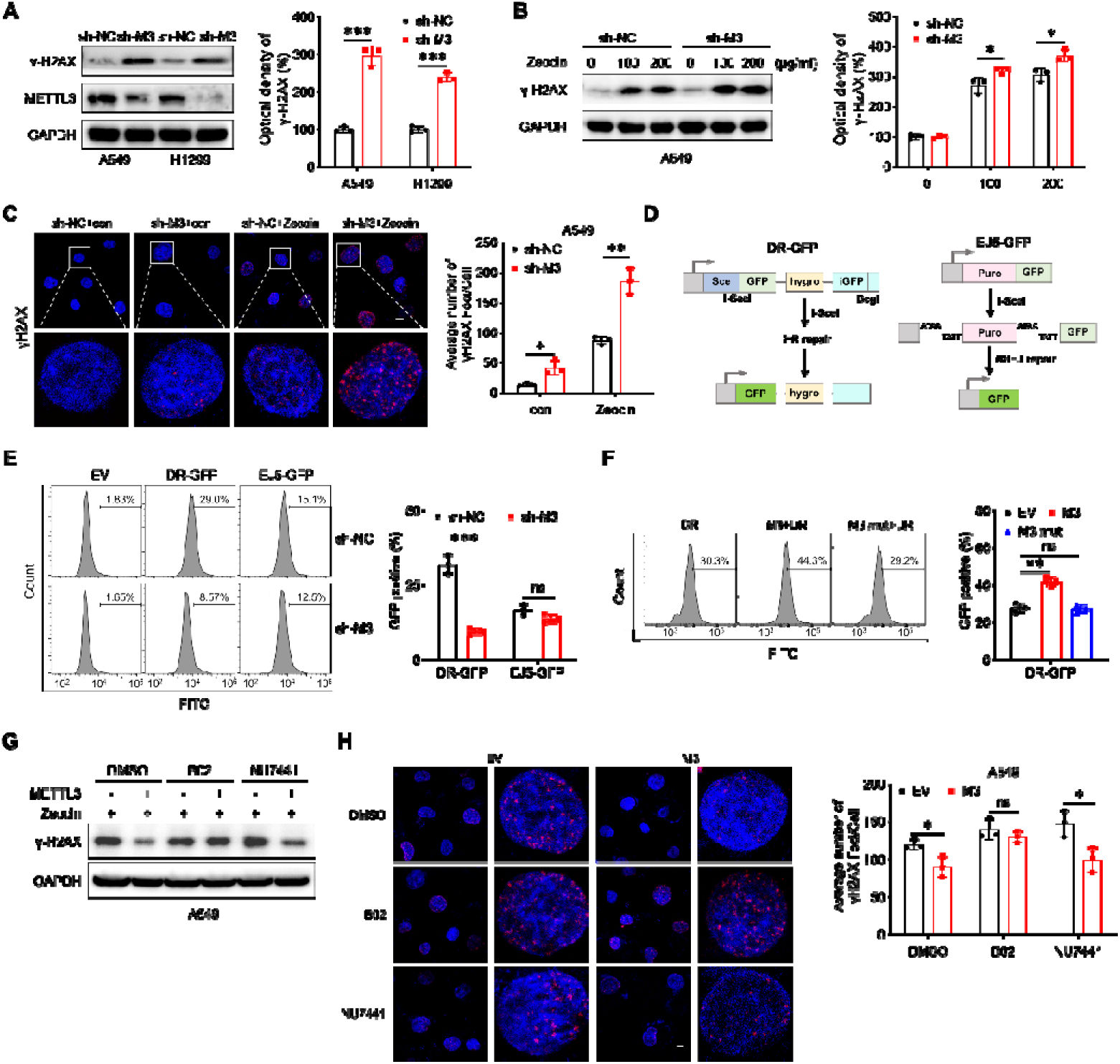
METTL3 regulates cytosolic DNA levels by modulating HR efficacy. (A) The protein expression of γH2AX in sh-control and sh-METTL3 A549 and H1299 cells (left) and quantitatively analyzed (right); (B) The protein expression of γH2AX in sh-control and sh-METTL3 A549 with Zeocin treatment (left) and quantitatively analyzed (right); (C) Representative confocal images of γH2AX in sh-control and sh-METTL3 A549 cells with or without Zeocin treatment, scale bar = 10 μm; (D) Schematic of the plasmid system for determining the frequency of HR- or NHEJ-mediated DSBR; (E) Quantification of the frequency of HR-mediated and NHEJ-mediated DSBR in sh-control and sh-METTL3 A549 cells; (F) Quantification of the frequency of HR-mediated DSBR in A549 cells transfected with vector control, METTL3 WT plasmid, METTL3 DA mutant plasmid for 24 h; (G) The protein expression of γH2AX in A549 cells transfected with vector control or METTL3 WT plasmid and then pretreated with B02 (a specific inhibitor of RAD51 and HR), NU7441 (a specific inhibitor of ligase IV and NHEJ), or DMSO and Zeocin treatment for 4 h; (H) Representative confocal images of γH2AX inA549 cells transfected with vector control or METTL3 WT plasmid and then pretreated with B02 (a specific inhibitor of RAD51 and HR), NU7441 (a specific inhibitor of ligase IV and NHEJ), or DMSO and Zeocin treatment for 4 h, scale bar = 10 μm. Data are presented as mean ± SD from three independent experiments. \**p*<0.05, \*\**p*<0.01, \*\*\**p*<0.001, ns, no significant, by Student’s *t* test between two groups and by one-way ANOVA followed by Bonferroni test for multiple comparison.

DSBs are primarily repaired by error-free homologous recombination repair (HR) or error-prone nonhomologous end joining (NHEJ)^[27]^. To investigate whether METTL3 influences one or both pathways, the HR and NHEJ reporter system were used (Figure 3 D). Results indicated that METTL3 deficiency impaired HR-mediated DSB repair (DSRB) but had no effect on NHEJ-mediated DSBR (Figures 3 E). Moreover, the over expression of METTL3, rather than METTL3 DA mutant, significantly promotes HR-mediated DSBR (Figures 3 F). It suggested that METTL3 can regulate the HR efficacy via a methyltransferase activity dependent manner. Furthermore, in the presence of B02, a specific inhibitor of RAD51 and HR^[28]^, overexpression of METTL3 failed to stimulate repair of Zeocin-induced DSBs. While NU7441, a specific inhibitor of ligase IV and NHEJ^[29]^, had no effect on METTL3-stimulated repair of Zeocin-induced DSBs (Figures 3 G and H). These findings suggest that knockdown of METTL3 accumulates cytosolic DNA by reducing the efficacy of HR-mediated DSBR.

### 4. MSH5 mediates METTL3-regualted HR efficacy and cytosolic DNA accumulation

To identify potential targets involved in m^6^A-regulated HR efficacy and cytosolic DNA accumulation, we performed mRNA-seq in sh-control and sh-*METTL3* A549 cells. Expression levels of 562 genes were found to be significantly changed with the up-regulation of 333 and down-regulation of 229 genes in sh-*METTL3* A549 cells (Figure S3 A, Supplementary Table 1). Among the 64 HR related genes (Supplementary Table 2), we identified the only candidate, mutS homolog 5 (MSH5), that overlapping among variated genes (greater than 2.0-fold variation (p < 0.05) between sh-control and sh-*METTL3* A549 cells, and m^6^A-modificated genes in A549 cells which was identified by m^6^A-seq (Supplementary Table 3) (Figure 4 A). RT-qPCR consistently showed that a reduction of METTL3 can decrease mRNA of *MSH5* in both A549 and H1299 cells (Figure 4 B). Consistently, western blot analysis showed that knockdown of METTL3 decreased protein expression of MSH5 in both A549 and H1299 cells (Figure 4 C).

**Figure 4.**
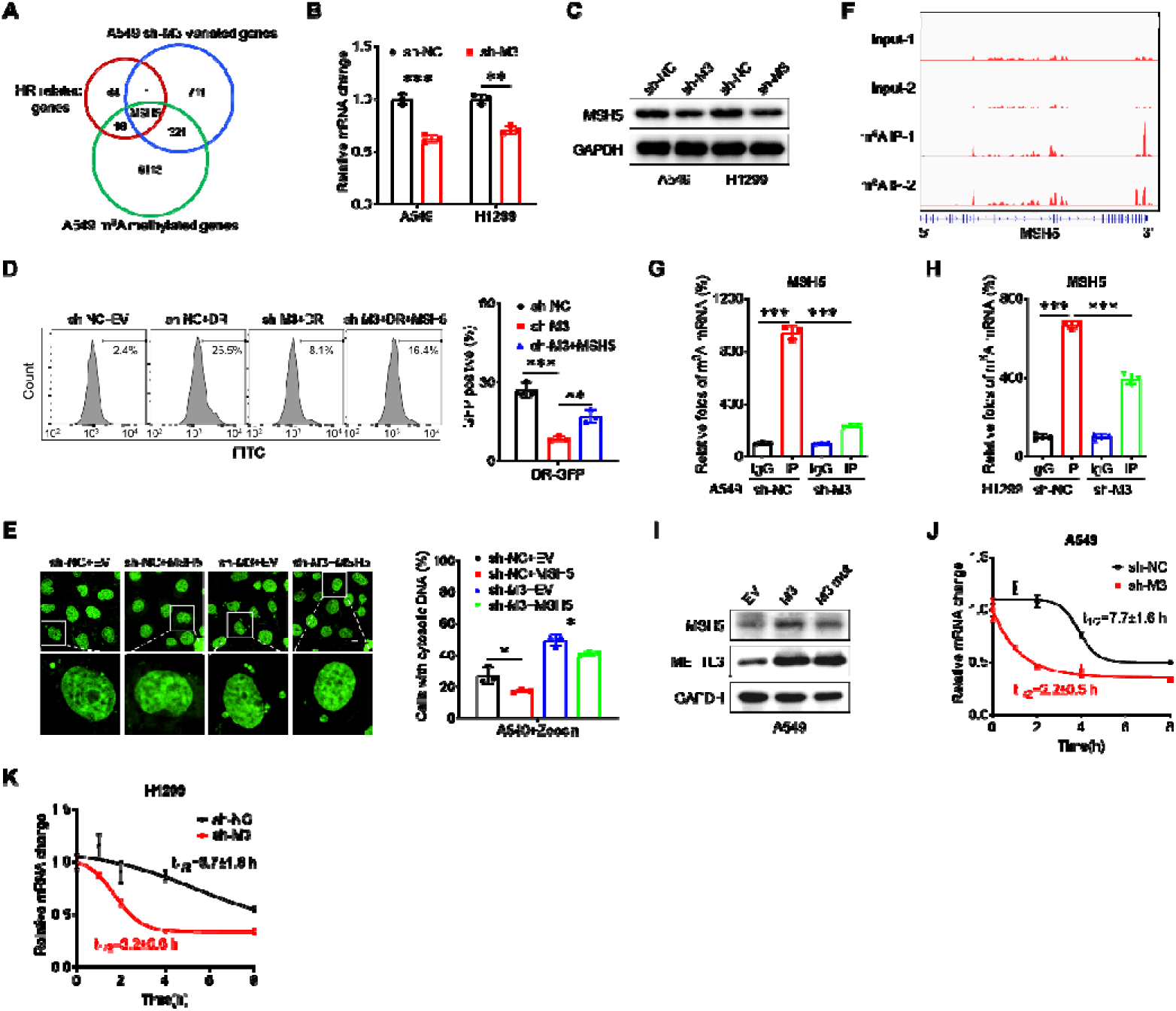
MSH5 mediates METTL3-regualted HR efficacy and cytosolic DNA accumulation. (A) Venn diagram shows substantial and significant overlap among HR related genes, variated genes between sh-METTL3 and sh-control A549 cells, and m^6^A-modificated genes in A549 cells; (B) The mRNA levels of *MSH5* in sh-control and sh-*METTL3* A549 and H1299 cells; (C) The protein expression of MSH5 in sh-control and sh-*METTL3* A549 and H1299 cells; (D) Quantification of the frequency of HR-mediated DSBR in A549 cells transfected with vector control or MSH5 plasmid for 24 h; (E) Representative confocal images of PicoGreen stain in sh-control and sh-METTL3 A549 cells transfected with vector control or MSH5 plasmid (left) and the percentages of cells displaying cytosolic DNA were measured (right), scale bar = 10 μm; (F) m^6^A peaks were enriched in *MSH5* mRNA from m^6^A RIP-seq data; (G) m^6^A RIP-qPCR analysis of *MSH5* in sh-control and sh-*METTL3* A549 cells; (H) m^6^A RIP-qPCR analysis of *MSH5* in sh-control and sh-*METTL3* H1299 cells; (I) The protein expression of MSH5 in A549 cells transfected with vector control, METTL3 WT plasmid, or METTL3 DA mutant plasmid for 24 h; (J) After treatment with Act-D for the indicated times, the mRNA levels of *MSH5* were checked in sh-control and sh-*METTL3* A549 cells; (K) After treatment with Act-D for the indicated times, the mRNA levels of *MSH5* were checked in sh-control and sh-*METTL3* H1299 cells. Data are presented as mean ± SD from three independent experiments. \**p*<0.05, \*\**p*<0.01, \*\*\**p*<0.001, ns, no significant, by Student’s *t* test between two groups and by one-way ANOVA followed by Bonferroni test for multiple comparison.

MSH5 encodes a member of the mutS family of proteins that are involved in DNA mismatch repair and meiotic recombination^[30]^. We therefore investigated its roles in m^6^A-regulated HR efficacy and cytosolic DNA accumulation of lung adenocarcinoma cells. Results showed that over expression of MSH5 (Figure S3 B) can reverse the down regulation of HR efficacy (Figure 4 D) and the increase of cytosolic DNA accumulation (Figure 4 E) in sh-*METTL3* A549 cells. It indicated that MSH5 mediates METTL3-regualted HR efficacy and cytosolic DNA accumulation in lung adenocarcinoma cells.

We further investigated whether m^6^A regulated the expression of MSH5 via direct methylation of mRNA. m^6^A-RIP-seq data showed that the CDS region of *MSH5* mRNA was modified by m^6^A (Figure 4 F). m^6^A-RIP-qPCR confirmed that a 10-fold enrichment of m^6^A antibody was observed in *MSH5* mRNA in A549 cells, and this enrichment significantly decreased in sh-*METTL3* A549 cells (Figure 4 G). Similar results were also observed in H1299 cells (Figure 4 H). Furthermore, the overexpression of WT METTL3, but not the METTL3 DA mutant, reversed the levels of MSH5 in A549 cells (Figure 4 I). It indicated that MSH5 was modified by m^6^A and that m^6^A positively regulates the expression of MSH5 in lung adenocarcinoma cells.

We further investigated the potential mechanisms involved in the m^6^A-regulated expression of MSH5. To determine whether METTL3 can modulate the transcription of MSH5, we performed a luciferase reporter assay by transfecting the promoter reporter gene plasmid pGL3-Basic-*MSH5*-luc into A549 cells. There was no significant difference in the luciferase activity of the MSH5 promoter between sh-control and sh-METTL3 lung adenocarcinoma cells (Figure S3 C), suggesting that METTL3 does not affect the transcription of *MSH5.* This was further confirmed by qRT-PCR analysis, which showed comparable levels of the precursor mRNA of MSH5 between sh-control and sh-METTL3 lung adenocarcinoma cells (Figure S3 D). Additionally, fractionation assay results indicated that there was no difference in the subcellular localization of *MSH5* mRNA in sh-control and sh-*METTL3* A549 cells (Figure S3 E). This indicated that m^6^A had no effect on the sub-cellular localization of *MSH5* mRNA.

However, knockdown of METTL3 (Figure 4 B) significantly decreased the mRNA expression of MSH5 in lung adenocarcinoma cells. Since m^6^A had no effect on promoter activity but regulates the mRNA expression of MSH5, we then tested its effect on mRNA stability. Our results showed that knockdown of METTL3 significantly decreased the mRNA stability of MSH5 in both A549 (Figure 4 J) and H1299 (Figure 4 K) cells. Therefore, the m^6^A-regulated expression of MSH5 should be down to the positive effects of m^6^A in mRNA stability of *MSH5* mRNA.

Regarding the translation efficiency of the endogenous MSH5 mRNA, which is defined as the ratio of protein production (MSH5/GAPDH) to mRNA abundance^[31]^, knockdown of METTL3 had no effects on the translation efficiency of MSH5 in A549 cells (Figure S3 F). To assess whether m^6^A can post-translationally regulate the expression of MSH5, both sh-control and sh-*METTL3* A549 cells were further treated with cycloheximide (CHX) to inhibit translation. Our data revealed that half-life of MSH5 protein had no significant difference between sh-Control and *sh-METTL3* cells (Figure S3 G). All these data indicated that METTL3 positively regulates the mRNA stability of *MSH5*, without affecting its transcription, nuclear export, translation efficiency, or protein stability.

### 5. METTL3 stabilized MSH5 mRNA via binding IGF2BP2 with methylation of A2521

We further investigated the methylation site and reader protein responsible for m^6^A-stablized *MSH5* mRNA. According to the m^6^A-seq results (Figure 4 F) and the prediction results of *MSH5* mRNA by use of m^6^A sites predictor SRAMP (http://www.cuilab.cn/sramp) (Figure S4 A), there were three GGAC motif in *MSH5* CDS was identified (Figure 5 A). m^6^A-RIP-PCR using of fragmented poly^+^ RNA indicated that the relative enrichment of CDS-3 of *MSH5* mRNA was much greater than that of two others in CDS in A549 cells (Figure 5 B). To investigate whether m^6^A methylated CDS-3 was involved in m^6^A-regulated mRNA stability of *MSH5* mRNA, we constructed CDS reporters containing wild type *MSH5* CDS behind the firefly luciferase reporter gene by use of the pmiR-GLO vector (Figure 5 C). The luciferase assay illustrated that the levels of F-Luc in sh-*METTL3* A549 cells were significantly decreased, which is due to the down regulation of *F-Luc* mRNA and not the translation efficiency (Figure 5 D). Further, the addition of *MSH5* CDS decreased the half-life of *F-Luc* mRNA in sh-*METTL3* A549 cells (Figure 5 E). It further confirmed that m^6^A-methylated CDS mediated METTL3-regulated mRNA stability of *MSH5*.

**Figure 5.**
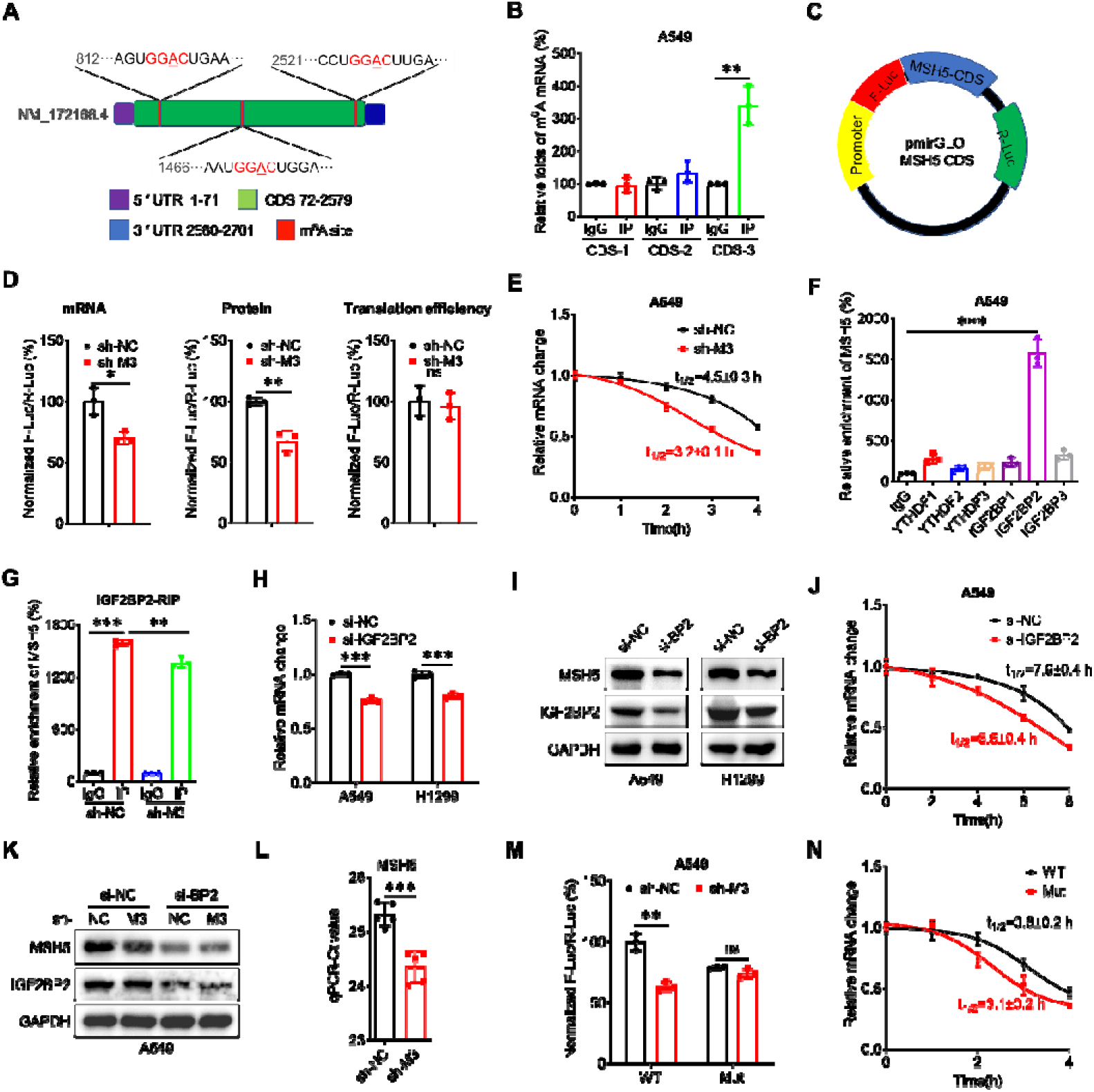
METTL3 stabilized MSH5 mRNA via binding IGF2BP2 with methylation of A2521. (A) Schematic representation of the three GGAC motifs in MSH5 CDS; (B) The m^6^A of the three GGAC motifs in MSH5 CDS in A549 cells were analyzed by m^6^A-RIP-qPCR using fragmented RNA; (C) Schematic representation of pmirGLO-*MSH5* CDS reporter; (D) The mRNA abundance, luciferase activity and translation efficiency of F-Luc in sh-control and sh-*METTL3* A549 cells transfected with pmirGLO-*MSH5* CDS reporter for 24 h; (E) After treatment with Act-D for the indicated times, the mRNA levels of *F-Luc* were checked in sh-control and sh-*METTL3* A549 cells transfected with pmirGLO-*MSH5* CDS reporter for 24 h; (F) RIP-qPCR analysis of *MSH5* mRNA in A549 cells by use of antibody of YTHDF1-3, and IGF2BP1-3; (G) IGF2BP2 RIP-qPCR analysis of *MSH5* mRNA in sh-control and sh-*METTL3* A549 cells; (H) The mRNA levels of MSH5 in A549 and H1299 cells transfected with si-NC or si-IGF2BP2 for 24 h; (I) The protein expression of MSH5 in A549 and H1299 cells transfected with si-NC or si-IGF2BP2 for 24 h; (J) After treatment with Act-D for the indicated times, the mRNA levels of *MSH5* were checked in A549 cells transfected with si-NC or si-IGF2BP2 for 24 h; (K) The protein expression of MSH5 in sh-control and sh-*METTL3* A549 cells transfected with si-NC or si-IGF2BP2 for 24 h; (L) The threshold cycle (Ct) of qPCR showing SELECT results for detecting m^6^A site in the potential m^6^A site of MSH5 in sh-control and sh-*METTL3* A549 cells; (M) The relative luciferase activity of F-Luc/R-Luc of pmirGLO-*MSH5* CDS WT, Mut reporter in sh-control and sh-*METTL3* A549 cells; (N) After treatment with Act-D for the indicated times, the mRNA levels of *F-Luc* were checked in sh-control and sh-*METTL3* A549 cells transfected with pmirGLO-*MSH5* CDS WT, Mut reporter for 24 h. Data are presented as mean ± SD from three independent experiments. \**p*<0.05, \*\**p*<0.01, \*\*\**p*<0.001, ns, no significant, by Student’s *t* test between two groups and by one-way ANOVA followed by Bonferroni test for multiple comparison.

The m^6^A binding proteins insulin-like growth factor 2 mRNA-binding proteins (IGF2BPs; including IGF2BP1/2/3), while not YTH domain family proteins YTHDF1/2/3, recognize and stabilize m^6^A-modified cellular RNAs ^[32]^. RIP-qPCR showed that IGF2BP2, not IGF2BP1/3 or YTHDF1/2/3, can bind with *MSH5* mRNA in A549 cells (Figure 5 F). Further, the binding between IGF2BP2 and *MSH5* mRNA was decreased in sh-*METTL3* A549 cells (Figure 5 G). Knockdown of IGF2BP2 was able to suppress the mRNA (Figure 5 H) and protein (Figure 5 I) expression of MSH5 in both A549 and H1299 cells. It was due to that knockdown of IGF2BP2 that there was suppression of the mRNA stability of MSH5 (Figure 5 J). Additionally, results showed that si-IGF2BP2 can suppress the expression of MSH5 and attenuate deletion of METTL3-suppressed expression of MSH5 in A549 cells (Figure 5 K). All these data suggested that IGF2BP2 mediated m^6^A-regulated expression of MSH5.

Our data suggested that **A2521** within CDS-3 might be the potential m^6^A site for *MSH5* mRNA. We further confirmed the potential site by the “SELECT” method^[33]^ in A549 cells. The results showed that knockdown of METTL3 decreased the methylation levels of A2521(Figure 5 L). While the nearby nucleotide A2526 without m^6^A modification had a significantly lower Ct value than that of A2521 (Figure S4 B). We then mutated A2521 within *MSH5* CDS to investigate its roles in m^6^A regulated mRNA stability of MSH5 (Figure S4 C). The mutation of A2521 in CDS, resulted in a down regulation of luciferase activity of F-Luc (Figure 5 M), while the mutation-induced down regulation of luciferase activity of F-Luc was reversed in sh-*METTL3* cells. Further, the mutation of CDS A2521 decreased the mRNA stability of F-Luc (Figure 5 N). All data confirmed that A2521 within *MSH5* CDS mediated m^6^A-regulated mRNA stability of *MSH5*.

### 6. METTL3 destabilized cGAS mRNA via binding YTHDF2 and A1545 methylation

Because knockdown of METTL3 inhibited cytoplasmic DNA- but not the cGAMP-induced expression of type I IFNs and inflammatory genes in lung adenocarcinoma cells (Figure 6 A and B), we reasoned that METTL3 promoted innate immune signaling at the level of cGAS. Consistently, qRT-PCR showed that knockdown of METTL3 increased the mRNA expression of cGAS in both A549 (Figure 6 C) and H1299 (Figure S5 A) cells. Western blot analysis showed that knockdown of METTL3 increased the protein expression of cGAS in lung adenocarcinoma cells (Figure 6 D).

**Figure 6.**
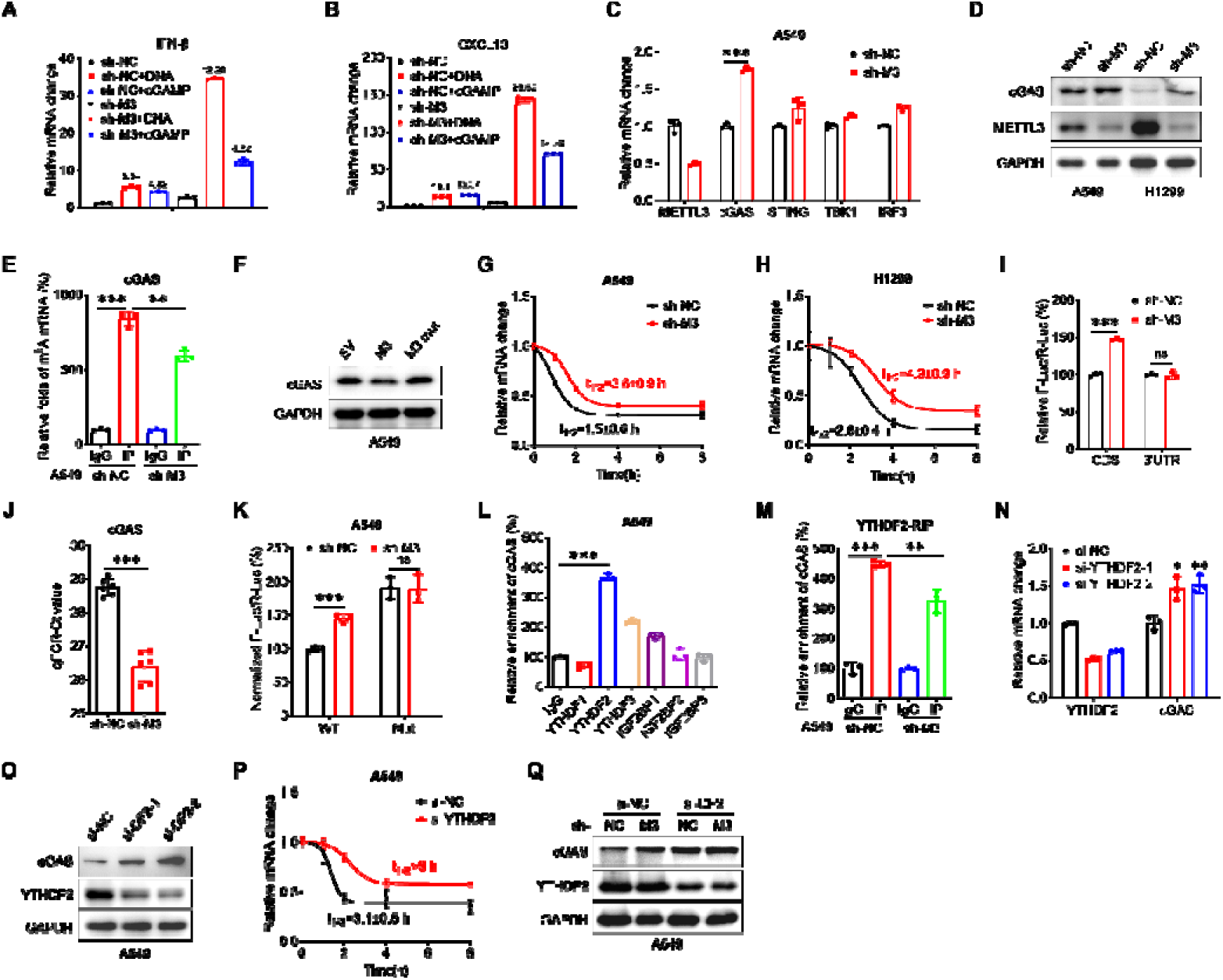
METTL3 destabilized cGAS mRNA via binding YTHDF2 and A1545 methylation. (A) The mRNA levels of IFN-β mRNA in sh-control and sh-METTL3 A549 cells treated with HT-DNA or cGAMP; (B) The mRNA levels of CXCL0 mRNA in sh-control and sh-METTL3 A549 cells treated with HT-DNA or cGAMP; (C) The mRNA expression of cGAS, STING, TBK1, IRF3 in sh-control and sh-METTL3 A549 cells; (D) The protein expression of cGAS in sh-control and sh-METTL3 A549 and H1299 cells; (E) m^6^A RIP-qPCR analysis of cGAS in sh-control and sh-*METTL3* A549 cells; (F) The protein expression of cGAS in A549 cells transfected with vector control, METTL3 WT plasmid, or METTL3 DA mutant plasmid for 24 h; (G) After treatment with Act-D for the indicated times, the mRNA levels of cGAS were checked in sh-control and sh-*METTL3* A549 cells; (H) After treatment with Act-D for the indicated times, the mRNA levels of cGAS were checked in sh-control and sh-*METTL3* H1299 cells; (I) The relative luciferase activity of F-Luc/R-Luc of pmirGLO-cGAS-CDS reporter or pmirGLO-cGAS-5’UTR reporter in sh-control and sh-*METTL3* A549 cells; (J) The threshold cycle (Ct) of qPCR showing SELECT results for detecting m^6^A site in the potential m^6^A site of cGAS in sh-control and sh-*METTL3* A549 cells; (K) The relative luciferase activity of F-Luc/R-Luc of pmirGLO-cGAS-CDS WT, Mut reporter in sh-control and sh-*METTL3* A549 cells; (L) RIP-qPCR analysis of cGAS mRNA in A549 cells by use of antibody of YTHDF1-3, and IGF2BP1-3; (M) YTHDF2 RIP-qPCR analysis of cGAS mRNA in sh-control and sh-*METTL3* A549 cells; (N) The mRNA levels of cGAS in A549 cells transfected with si-NC or si-YTHDF2-1/2 for 24 h; (O) The protein expression of cGAS in A549 cells transfected with si-NC or si-YTHDF2-1/2 for 24 h; (P) After treatment with Act-D for the indicated times, the mRNA levels of cGAS were checked in sh-control and sh-*METTL3* A549 cells transfected with si-NC or si-YTHDF2 for 24 h; (Q) The protein expression of cGAS in sh-control and sh-*METTL3* A549 cells transfected with si-NC or si-YTHDF2 for 24 h; Data are presented as mean ± SD from three independent experiments. \**p*<0.05, \*\**p*<0.01, \*\*\**p*<0.001, ns, no significant, by Student’s *t* test between two groups and by one-way ANOVA followed by Bonferroni test for multiple comparison.

We further investigated whether m^6^A regulated expression of cGAS via directly methylate its mRNA. m^6^A-RIP-qPCR confirmed an 8-fold m^6^A antibody enriched *cGAS* mRNA in A549 cells, while this enrichment significantly decreased in sh-*METTL3* A549 cells (Figure 6 E). Similar results were also observed in H1299 cells (Figure S5 B). It indicated that *cGAS* mRNA was m^6^A methylated. Further, the transfections of WT METTL3, while not catalytically inactive METTL3 mutant DA (D395A), decreased the expression of cGAS in A549 cells (Figure 6 F). Additionally, STM2457 significantly increased the protein expression of cGAS in A549 cells (Figure S5 C). It indicated that METTL3 can positively regulate the expression of cGAS in lung adenocarcinoma cells via an m^6^A dependent manner.

We further investigated the mechanisms for m^6^A-regulated expression of cGAS. Data suggest the transcription (Figure S5 D), sub-cellular localization (Figure S5 E), translation efficiency (Figure S5 F) of *cGAS* mRNA and half-life of cGAS protein (Figure S5 G) in sh-control and sh-*METTL3* cells were comparable. We then tested its effect on mRNA stability. The results showed that knockdown of METTL3 significantly increased the mRNA stability of *cGAS* in both A549 (Figure 6 G) and H1299 (Figure 6 H) cells. All these data indicated that METTL3 negatively regulated mRNA stability of *cGAS*, while had no effect on its transcription, nuclear export, translation efficiency, or protein stability.

We further investigated the methylation site and reader protein responsible for m^6^A-stablized *cGAS* mRNA. As shown in Figure S5 H, three GGAC motif in *cGAS* mRNA was identified by the m^6^A sites predictor. To investigate whether m^6^A-methylated CDS or 5’UTR were involved in m^6^A-regulated stability of *cGAS* mRNA, we constructed CDS and 5’UTR reporters by inserting the wild-type cGAS CDS or 3’UTR downstream of the firefly luciferase reporter gene by use of pmiR-GLO vector (Figure S6 I). The luciferase assay revealed a significant increase in the luciferase expression of F-Luc of pmiR-GLO-cGAS-CDS reporter rather than pmiR-GLO-cGAS-3’UTR reporter in sh-*METTL3* A549 cells (Figure 6 I). The m^6^A methylation of **A1545** within the GGAC motif in *cGAS* CDS was confirmed by the “SELECT” method in A549 cells, whilst knockdown of METTL3 can decrease the methylation levels of A1545 (Figure 6 J). While the nearby nucleotide A1542 without m^6^A modification had significant lower Ct value than that of A1545 (Figure S5 J). The mutation of A1545 in CDS resulted in an increase of F-Luc activity, which was abolished in sh-*METTL3* cells (Figure 6 K). All these data confirmed that methylation of **A1545** within *cGAS* CDS was involved in m^6^A regulated cGAS.

We further checked the reader protein responsible for m^6^A-regualted expression of cGAS. RIP-qPCR showed that YTHDF2, and not IGF2BP1/2/3 or YTHDF1/3 can bind with *cGAS* mRNA (Figure 6 L). Further, the binding between YTHDF2 and *cGAS* mRNA was decreased in sh-*METTL3* A549 cells (Figure 6 M). Knockdown of YTHDF2 raised the mRNA (Figure 6 N) and protein (Figure 6 O) expression of cGAS in A549 cells. It was due to that knockdown of YTHDF2 increased mRNA stability of cGAS (Figure 6 P). Additionally, results showed that si-YTHDF2 can increase the expression of cGAS and attenuate deletion of METTL3- increased expression of cGAS in A549 cells (Figure 6 Q). All data suggested that YTHDF2 mediated m^6^A-regulated expression of cGAS.

### 7. METTL3 inhibition enhances immunotherapy and suppresses cancer progression in lung adenocarcinoma

To investigate the therapeutic efficacy of combining METTL3 inhibition with immunotherapy, we established syngeneic tumor models using sh-control or sh-METTL3 LLC cells, which were treated with or without anti-PD-1 antibody. Results revealed that knockdown of METTL3 substantially enhanced the efficacy of PD-1 antibody therapy, leading to a marked reduction in tumor volume and weight (Figure 7 A-C). Subsequently, flow cytometry analysis demonstrated that METTL3 inhibition in combination with PD-1 antibody significantly increase the proportion of CD8^+^ T cells within the tumor tissue (Figure 7 D). These findings collectively indicate that METTL3 inhibition improves tumor immune infiltration and enhances the efficacy of immunotherapy.

**Figure 7.**
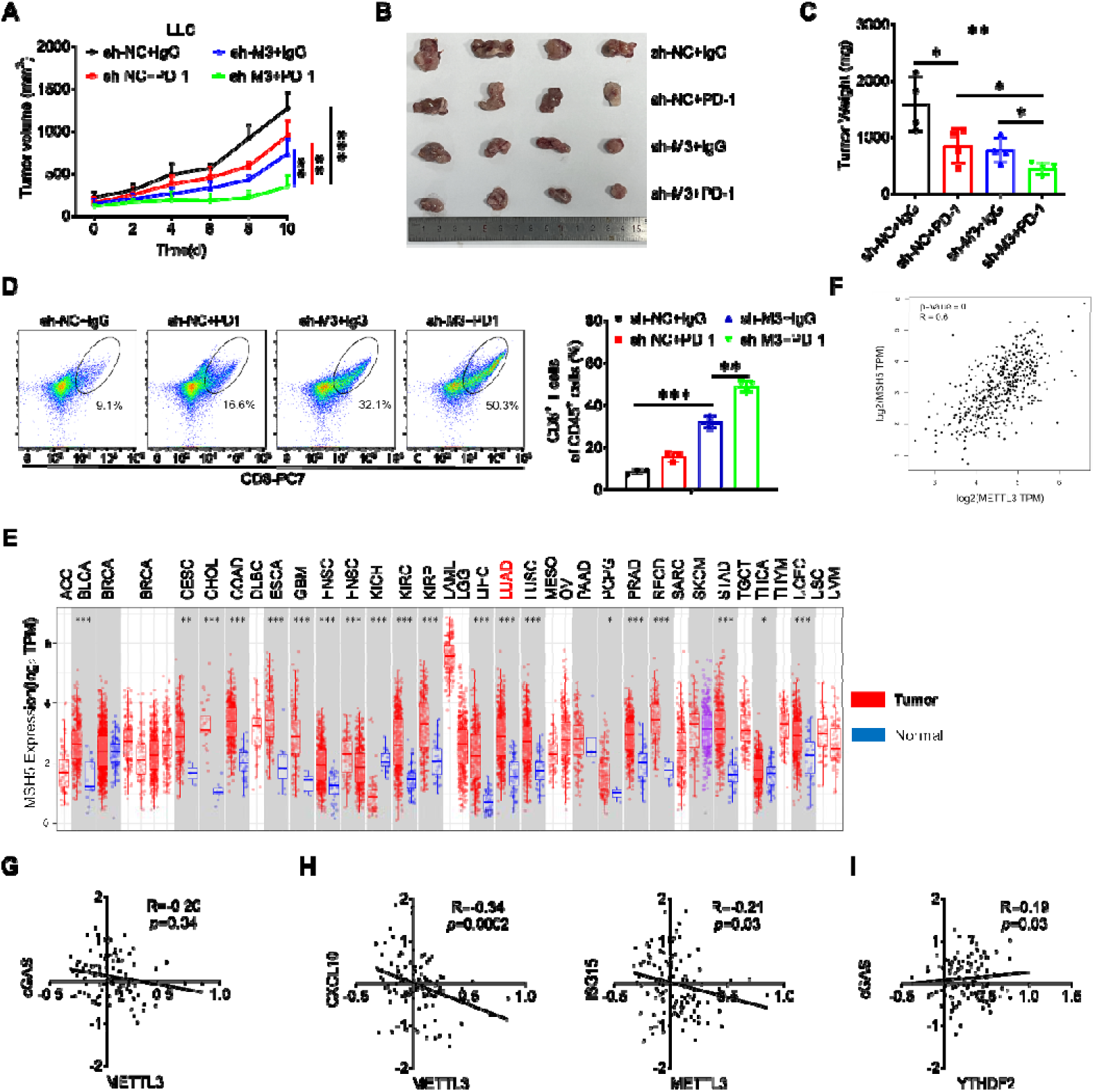
METTL3 inhibition enhances immunotherapy and suppresses cancer progression in lung adenocarcinoma. (A) The tumor growth curves of syngeneic tumor models using sh-control or sh-METTL3 LLC cells treated with or without anti-PD-1 antibody; (B) The tumor volume of syngeneic tumor models using sh-control or sh-METTL3 LLC cells treated with or without anti-PD-1 antibody; (C) The tumor weight of of syngeneic tumor models using sh-control or sh-METTL3 LLC cells treated with or without anti-PD-1 antibody; (D) The percentages of CD8^+^ T cell in CD45^+^ cell in the tumor tissues taken from mice with sh-control or sh-METTL3 LLC cells syngeneic tumor treated with or without anti-PD-1 antibody; (E) The expression of MSH5 across different types of cancers based on TCGA database; (F) The correlation between the expression of METTL3 and MSH5 in lung adenocarcinoma based on TCGA database; (G) The correlation between the expression of METTL3 and cGAS in lung adenocarcinoma from CPTAC database; (H) The correlation between the expression of METTL3 and CXCL10, ISG15 in lung adenocarcinoma from CPTAC database; (I) The correlation between the expression of YTHDF2 and cGAS in lung adenocarcinoma from CPTAC database. Data are presented as mean ± SD from three independent experiments. \**p*<0.05, \*\**p*<0.01, \*\*\**p*<0.001, ns, no significant, by Student’s *t* test between two groups and by one-way ANOVA followed by Bonferroni test for multiple comparison.

At this point, we investigated the potential connection between METTL3 inhibition and lung adenocarcinoma development using clinical data from databases. The data form TCGA indicated an upregulation in the expression of *MSH5* (Figure 7 E) in LUAD tumor tissues compared to normal tissues. Furthermore, data from GEIPA demonstrated a positive correlation between METTL3 and MSH5 (Figure 7 F) in LUAD cancer patients. Consistently, the protein expression of METTL3 was negatively correlated with cGAS (Figure 7 G) and ISG CXCL10, ISG15 (Figure 7 H) in LUAD cancer patients from the CPTAC database. Moreover, the expression of YTHDF2 exhibited a positive correlation with the cGAS (Figure 7 I) in LUAD cancer patients. Together, these data suggested that METTL3 inhibition-activated cGAS/STING axis suppressed LUAD progression.

### 8. METTL3 inhibition and PARP inhibitor combined enhances the anti-tumor functions of PD-1 antibody

Poly (ADP ribose) polymerase 1 (PARP1) is responsible for initiating the repair of single-strand break (SSB), which can potentially evolve into detrimental DSBs^[34]^. Recent studies showed that PARP inhibitors can induce micronuclei formation in cancer cell lines and trigger ISGs in a cGAS- and STING-dependent manner^[35, 36]^. The above research indicated that METTL3 deficiency reduces the homologous recombination repair efficacy, thereby activating the cGAS/STING-mediated anti-tumor immunity. We therefore explored whether METTL3 regulates chemotherapeutic response to PARP inhibitors in lung adenocarcinoma cells. Cell viability assay revealed that knockdown of METTL3 significantly increased the sensitivity of A549 and H1299 cells to Olaparib, a PARP inhibitor (Figure 8 A). Further, colony formation assay showed that METTL3 knockdown markedly inhibited colonization of both A549 and H1299 cells (Figure 8 B). In contrast, overexpression of METTL3 decreased the sensitivity and enhanced the colony formation of A549 cells to the treatment of Olaparib (Figure 8 C and D). These data suggested that loss of METTL3 sensitizes tumor cell to PARP inhibitors and impairs oncogenic transformation.

**Figure 8.**
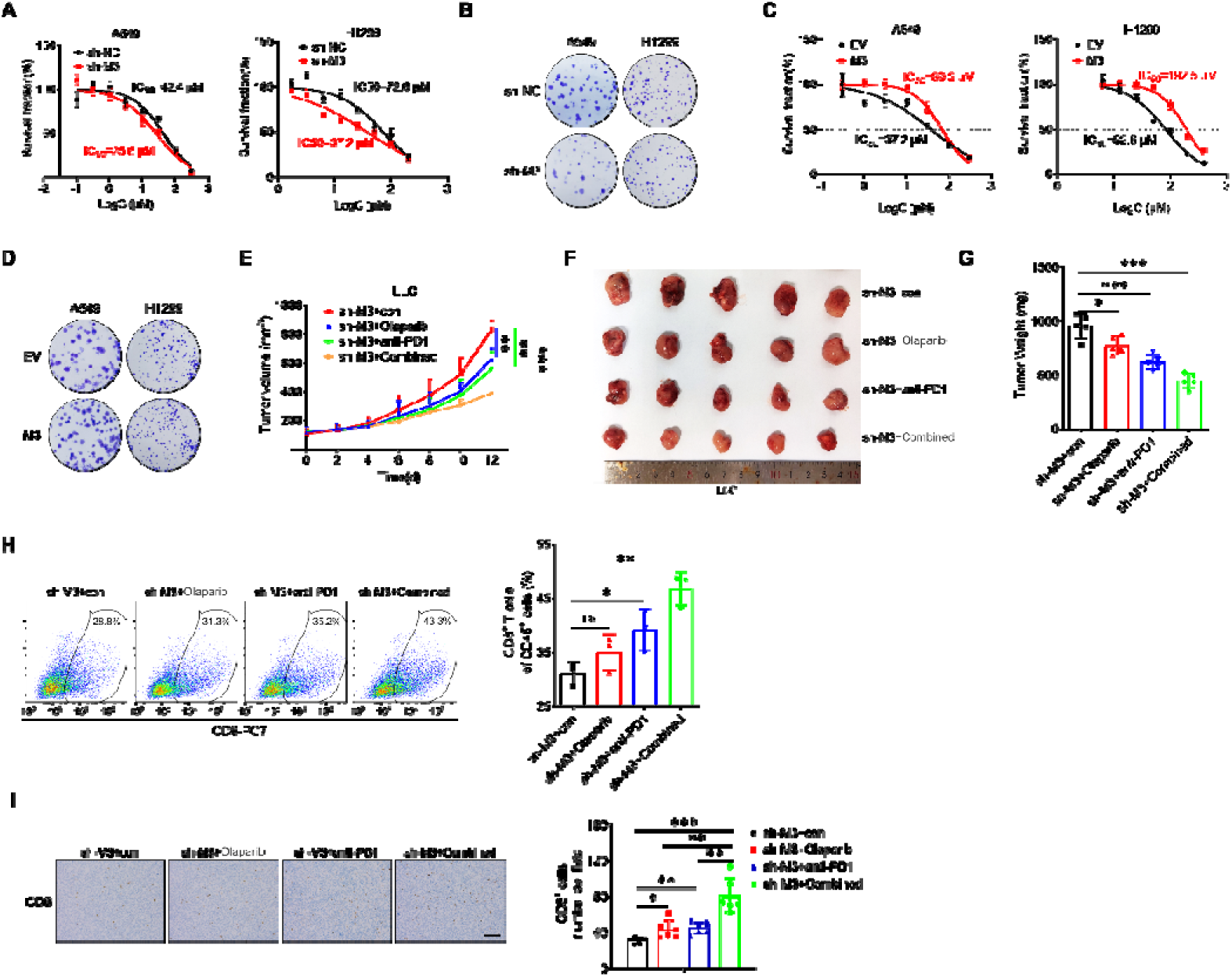
METTL3 inhibition and PARP inhibitor combined enhances the anti-tumor functions of PD-1 antibody. (A) The sensitivity of sh-control or sh-METTL3 A549 and H1299 cells to Olaparib; (B) The colony formation capacity of sh-control or sh-METTL3 A549 and H1299 cells treated with Olaparib; (C) The sensitivity of A549 and H1299 cells transfected with vector control, METTL3 WT plasmid for 24 h to Olaparib; (D) The colony formation capacity of A549 and H1299 cells transfected with vector control, METTL3 WT plasmid for 24 h treated with Olaparib; (E) The tumor growth curves of syngeneic tumor models using sh-*METTL3* LLC cells with Olaparib, anti-PD-1 antibody, or a combination of both treatment; (F) The tumor volume of using sh-*METTL3* LLC cells with Olaparib, anti-PD-1 antibody, or a combination of both treatment at the end of the experiment; (G) The tumor weight of using sh-*METTL3* LLC cells with Olaparib, anti-PD-1 antibody, or a combination of both treatment; (H) The percentages of CD8^+^ T cell in CD45^+^ cell in the tumor tissues taken from mice with sh-*METTL3* LLC syngeneic tumor with Olaparib, anti-PD-1 antibody, or a combination of both treatment; (I) IHC (CD8)-stained paraffin-embedded sections obtained from sh-*METTL3* LLC syngeneic tumor with Olaparib, anti-PD-1 antibody, or a combination of both treatment, the scale bar is 100 μm. Data are presented as mean ± SD from three independent experiments. \**p*<0.05, \*\**p*<0.01, \*\*\**p*<0.001, ns, no significant, by Student’s *t* test between two groups and by one-way ANOVA followed by Bonferroni test for multiple comparison.

Thus, we treated the sh-METTL3 LLC syngeneic tumor models with Olaparib, anti-PD-1 antibody, or a combination of both and monitored tumor growth (Figure 8 E). Notably, PARP inhibitor or anti-PD-1 antibody alone restricted tumor growth and tumor weight, while the combination of PARP inhibitor and anti-PD-1 antibody substantially offered significantly improved therapeutic effects compared with single treatment (Figure 8 F and G). Additionally, flow cytometry analysis revealed that each treatment increased intratumoral infiltration of CD8^+^ T cells and this effect was enhanced by the combined treatment (Figure 8 H). IHC analysis demonstrated that CD8 expression levels were further enhanced by the combined treatment (Figure 8 I). Collectively, our study demonstrates that targeting METTL3 and PARP inhibitor combined enhances the anti-tumor functions of PD-1 antibody.

## Discussion

Immunotherapies targeting PD-1/PD-L1-immune checkpoints have achieved unprecedented success in treating various types of cancers. However, immune suppression and treatment resistance have seriously limited its clinical efficacy in lung adenocarcinoma^[37]^. There is an urgent need to explore new strategies to enhance immunotherapies in lung adenocarcinoma. Our present study revealed that METTL3 inhibition-activated cGAS/STING axis enhances immunotherapy and suppresses cancer progression in lung adenocarcinoma via regulation of HR repair efficacy and the expression of cGAS. Mechanistically, IGF2BP2 bound with the A2521 in MSH5 CDS increase its’ mRNA stability, YTHDF2 bound with the A1545 in cGAS CDS decrease its’ mRNA stability, respectively. Collectively, METTL3 inhibition-activated cGAS/STING axis enhances immunotherapy and PARP inhibitor sensitivity via induction of cytosolic DNA and cGAS expression, which in turn regulate cancer progression in lung carcinoma (Figure 9). Our results describe the potential roles of METTL3 in immunotherapy, and also create the possibility to develop therapeutic strategies against LUAD progression by targeting METTL3/ cGAS axis.

**Figure 9.**
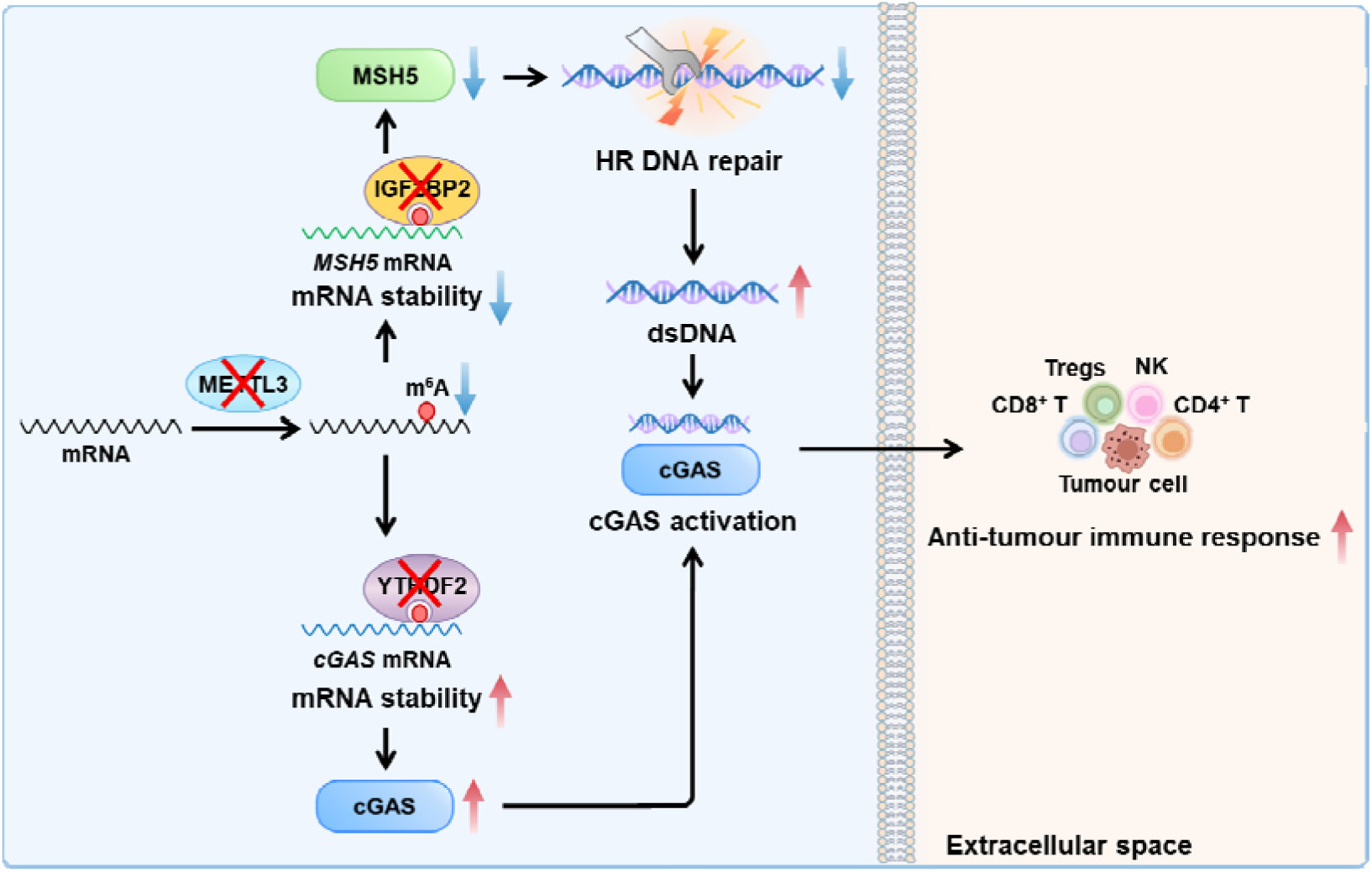

Previous research has elucidated the pivotal role of m^6^A modification in modulating the physiological behaviors of immune cells, such as infiltration, survival, differentiation, or polarization, in the tumor microenvironment^[38]^. The impact of m^6^A on immunotherapy efficacy is profound, as it can significantly influence the function of immune cells. For instance, the absence of YTHDF1 enhances the translation efficiency of lysosomal histone proteases in dendritic cells (DCs), thereby potentiating the anti-tumor activity of CD8^+^ T cells^[39]^. Additionally, Cluster analysis based on m^6^A features has uncovered a close correlation between m^6^A modification and the formation of the TME, indicating that targeting m^6^A in tumor cells may represent a promising strategy to regulate the TME and enhance the efficacy of tumor immunotherapy^[40]^. Herein, we found that Knockdown of METTL3 promotes the innate immune response and activates the cGAS-STING pathway. One recent study consistently showed that METTL3 inhibition has been shown to stimulate an endogenous interferon response in cells through the generation of double-stranded RNA, thereby enhancing anti-tumor effects^[41]^. Our data and previous studies confirmed the regulatory effects of m^6^A on innate immune response.

We identified that METTL3 regulates cytosolic DNA levels by modulating HR repair efficacy. Mechanism research indicated that MSH5 mediates METTL3-regualted HR efficacy and cytosolic DNA accumulation. MSH5 encodes a member of the mutS family of proteins that are involved in DNA mismatch repair and meiotic recombination^[30]^ and is also considered a vital component in tumor progression^[42, 43]^. Our data suggests that m^6^A/MSH5-regulated HR repair efficacy is a potential target for lung adenocarcinoma therapy. We further identified that m^6^A methylation of A1545 in cGAS decrease its mRNA stability and is involved in m^6^A-regualted cGAS pathway activation of lung adenocarcinoma cells. As the key sensor of the cytosolic accumulated DNA, cGAS emerges as an intriguing candidate for targeted cancer therapy^[44, 45]^. Post-transcriptional regulation of cGAS/STING signaling pathway has been extensively studied^[16]^, Our present study revealed another layer of regulation factors for cGAS expression.

Interestingly, the m^6^a modification sites on the MSH5 and cGAS mRNA are both located within their CDS regions, yet exhibit contrasting effects. The A2521 in MSH5 CDS binding with IGF2BP2 enhances its mRNA stability, whereas the A1545 in cGAS CDS binding with YTHDF2 reduces its mRNA stability. This divergence can be ascribed to the dissimilarity in the binding proteins, which act as the functional executors of the m^6^A modifications^[46]^.

Overall, our study sheds light on a novel relationship between innate immune response and m^6^A methylation. Specifically, METTL3 inhibition activates cGAS/STING-mediated anti-tumor immunity in lung carcinoma via HR efficacy and upregulation of cGAS. Given the numerous genes involved in innate immune response, it is plausible that m^6^A modification indirectly regulates innate immune response by affecting other genes. Our study suggested that METTL3 inhibition activates cGAS/STING-mediated anti-tumor immunity, which has expanded our understanding of such interplays that are essential for therapeutic application.

## Acknowledgements

We thank Prof Myan Cai at the Cancer Center of Sun Yat-sen University for plasmid donation, and Prof Feng Liu at the School of Life Sciences, Sun Yat-sen University, for experimental skills and instrumental help.

## Author contributions

HS Wang and JW Zhou designed and initiated the study;

JW Zhou, JX He and YQ Lu performed experiments;

C Yi, X Chang, LJ Tao, K Zhong, HS Zhang, JX Li, and ZJ Chen helped design the study and interpret the data;

JW Zhou and HS Wang wrote the paper.

## Data Availability Statement

The data supporting the findings of this study are available within the article and its supplementary materials.

## Availability of supplementary data

Nine figures and three tables are attached.

