## Supplementary data for "METTL3 inhibition-activated cGAS/STING axis enhances immunotherapy and PARP inhibitor sensitivity of lung carcinoma"

**Zhou et al**

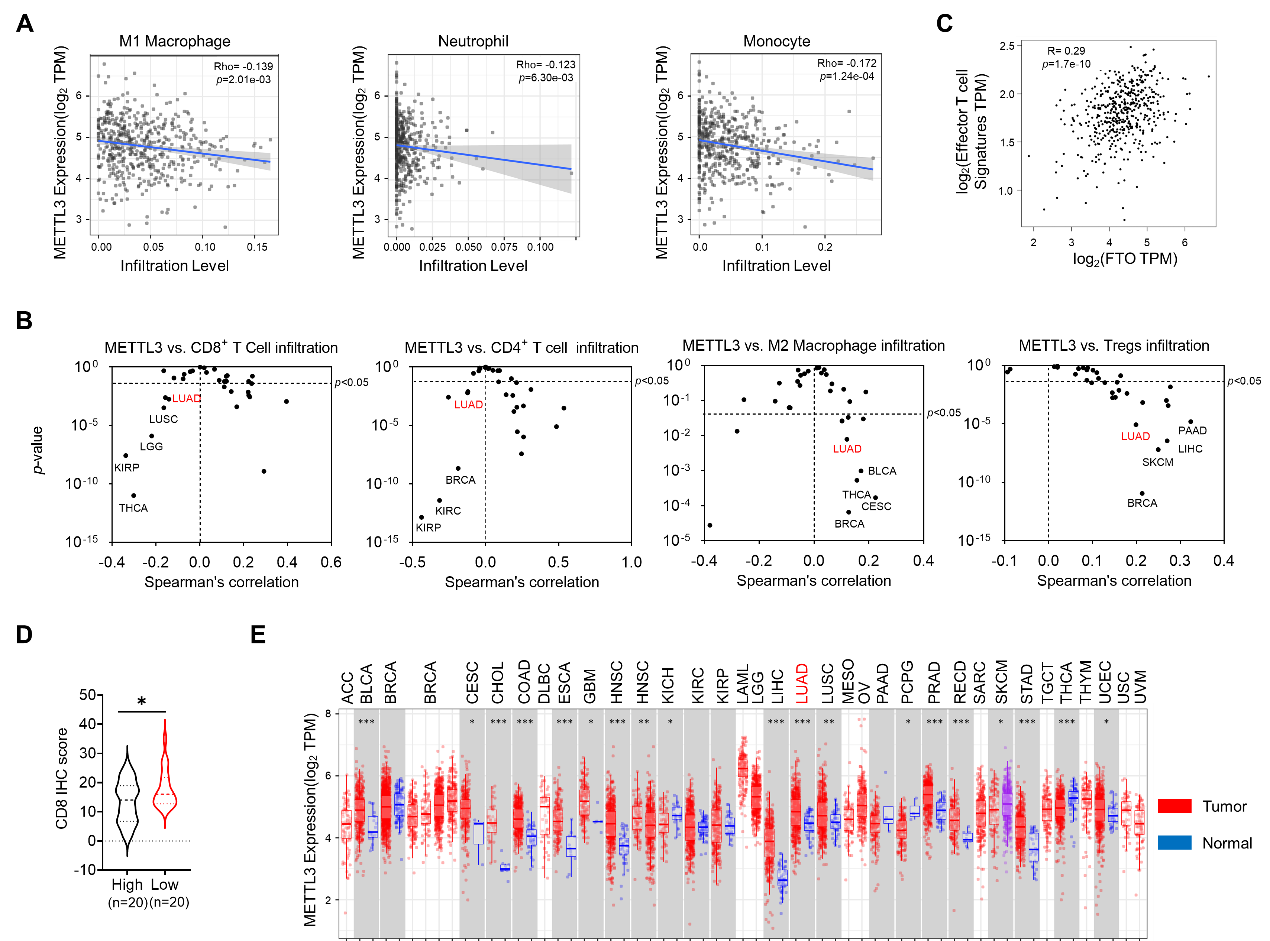

**Supplementary Figure 1 METTL3 expression is correlated with immune cell infiltration and cancer progression.**

(A) The correlation between the expression of METTL3 and infiltrating level of M1 macrophages, monocytes, or neutrophils in lung adenocarcinoma based on TIMER platform;
(B) The correlation between the expression of METTL3 and infiltrating level of CD8^+^ T cells, CD4^+^ T cells, M2 macrophages or Tregs across different types of cancers based on TIMER platform;
(C) The correlation between the expression of FTO and effector T cell markers (including CX3CR1, FGFBP2, FCGR3A) in lung adenocarcinoma based on TCGA database;
(D) The infiltrating level of CD8^+^ T cells from high or low IHC scores of METTL3 in lung adenocarcinoma tissues；
(E) The expression of METTL3 across different types of cancers based on TCGA database.

Data are presented as mean ± SD from three independent experiments. **p*<0.05, ***p*<0.01, ****p*<0.001, ns, no significant, by Student’s *t* test between two groups and by one-way ANOVA followed by Bonferroni test for multiple comparison.

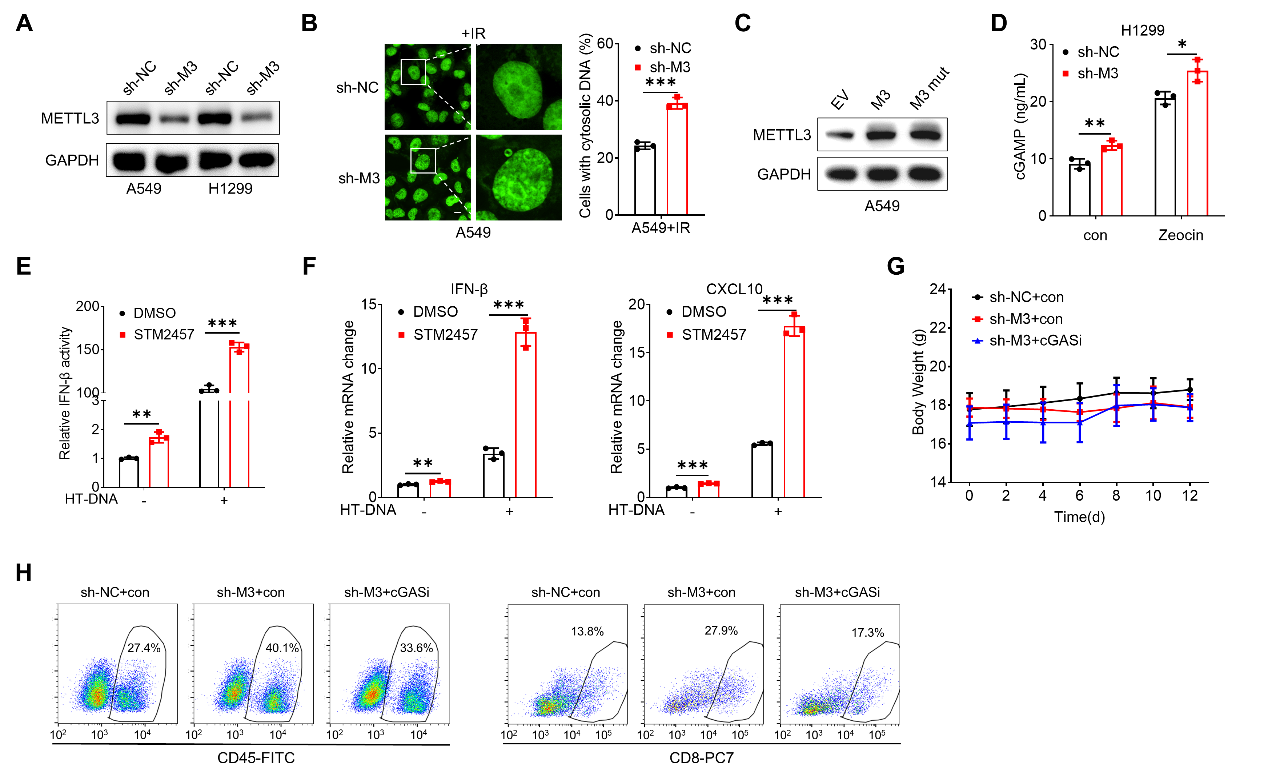

**Supplementary Figure 2 Knockdown of METTL3 promotes the innate immune response and activates the cGAS-STING pathway.**

(A) The protein expression of METTL3 in sh-*METTL3* A549, sh-*METTL3* H1299 cells and their corresponding control cells;
(B) Representative confocal images of PicoGreen stain in sh-control and sh-METTL3 A549 cells with IR treatment (left) and the percentages of cells displaying cytosolic DNA were measured (right), scale bar = 10 μm;
(C) The protein expression of METTL3 in A549 cells transfected with vector control, METTL3 WT plasmid, METTL3 DA mutant plasmid for 24 h;
(D) Intracellular cGAMP levels in sh-control and sh-METTL3 H1299 cells with and without Zeocin treatment;
(E) The mRNA levels of IFN-β, CXCL0 mRNA in A549 cells pretreated with STM2457 for 24 h and then treated with HT-DNA for 4 h;
(F) The IFN-β promoter activities in sh-control and sh-METTL3 A549 cells treated with HT-DNA;
(G) The body weight of mice with sh-control, sh-*METTL3* LLC xenografts with or without G140 treatment;
(H) Flow cytometry analysis of the percentages of CD45^+^ cell and the percentages of CD8^+^ T cell in CD45^+^ cell in the tumor tissues taken from mice with sh-control, sh-*METTL3* LLC xenografts with or without G140 treatment.

Data are presented as mean ± SD from three independent experiments. **p*<0.05, ***p*<0.01, ****p*<0.001, ns, no significant, by Student’s *t* test between two groups and by one-way ANOVA followed by Bonferroni test for multiple comparison.

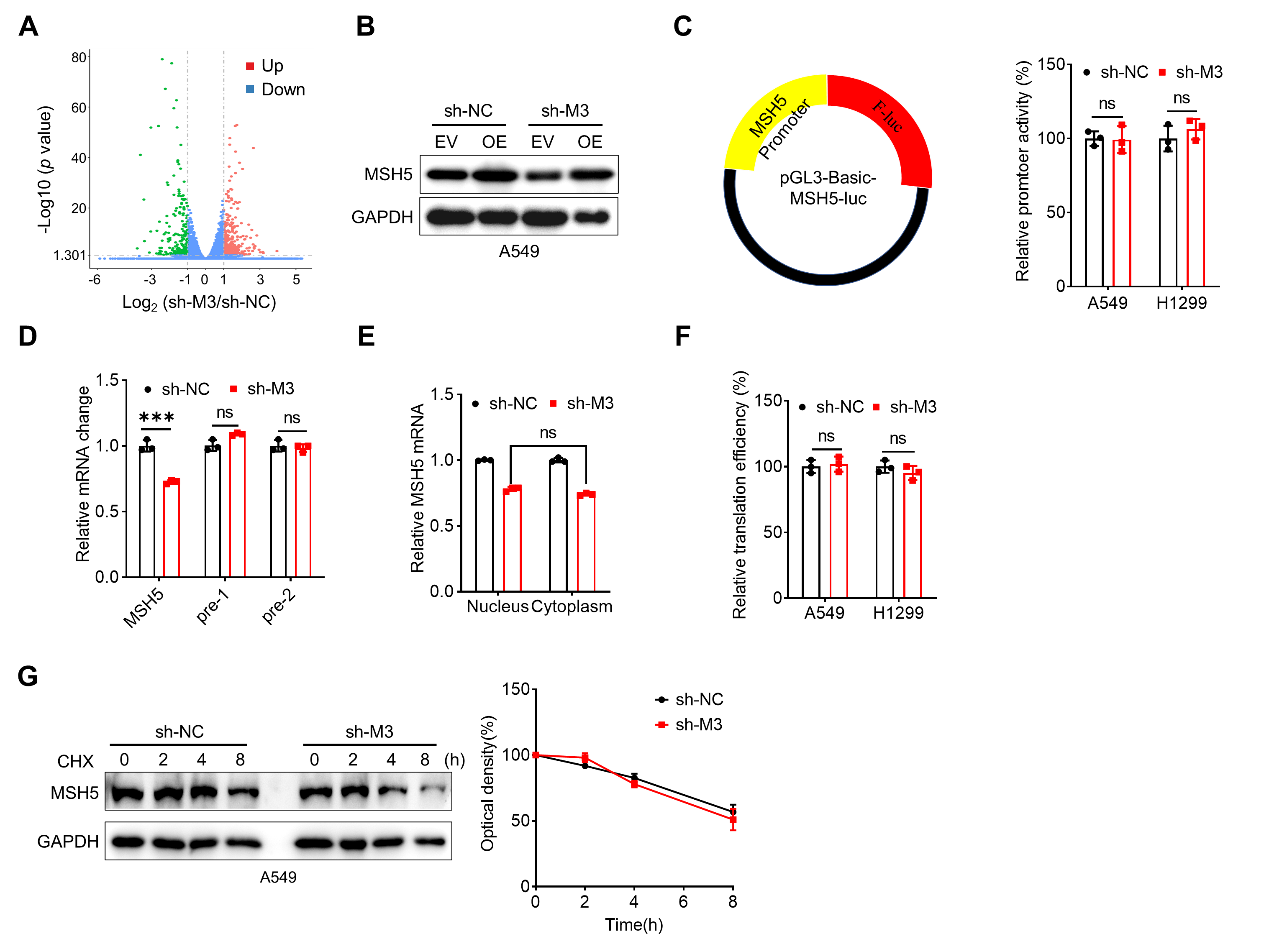

**Supplementary Figure 3 MSH5 mediates METTL3-regualted HR efficacy and cytosolic DNA accumulation.**

(A) Volcano plots to determine different genes in sh-*METTL3* cells as compared with that in A549 cells, each dot represents a gene;
(B) The protein expression of MSH5 in A549 cells transfected with vector control or MSH5 plasmid for 24 h;
(C) Cells were transfected with pGL3-Basic-*MSH5*-luc reporter and pRL-TK plasmid for 24 h, the promoter activities were presented as the ratios of the reporter normalized to pRL-TK plasmid;
(D) The levels of precursor *MSH5* mRNA in sh-control and sh-*METTL3* A549 cells;
(E) The relative levels of nuclear versus cytoplasmic *MSH5* mRNA in sh-control and sh-*METTL3* A549 cells;
(F) The translation efficiency of endogenous MSH5 in sh-control and sh-*METTL3* A549 cells;
(G) Cells were treated with 10 μg/ml CHX for the indicated time periods, the protein expression of MSH5 was detected by western blot analysis (left) and quantitatively analyzed (right).

Data are presented as mean ± SD from three independent experiments. **p*<0.05, ***p*<0.01, ****p*<0.001, ns, no significant, by Student’s *t* test between two groups and by one-way ANOVA followed by Bonferroni test for multiple comparison.

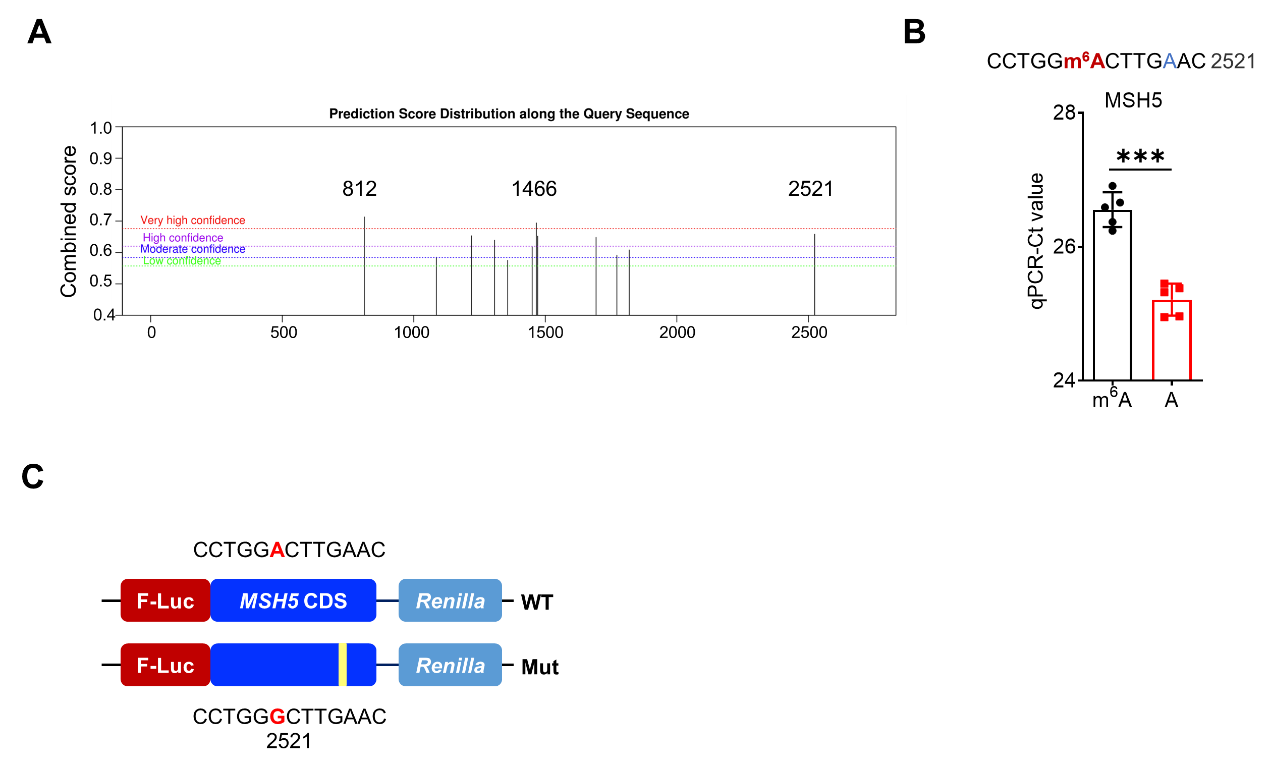

**Supplementary Figure 4 METTL3 stabilized MSH5 mRNA via binding IGF2BP2 with methylation of A2521.**

(A) The predicted m^6^A peaks in *MSH5* mRNA from the m^6^A sites predictor SRAMP;
(B) The threshold cycle (Ct) of qPCR showing SELECT results for detecting m^6^A site in the potential m^6^A site (A2521) and negative A site (A2526) of MSH5 in A549 cells;
(C) Schematic representation of mutation in CDS to investigate the m^6^A roles on MSH5 expression.

Data are presented as mean ± SD from three independent experiments. **p*<0.05, ***p*<0.01, ****p*<0.001, ns, no significant, by Student’s *t* test between two groups and by one-way ANOVA followed by Bonferroni test for multiple comparison.

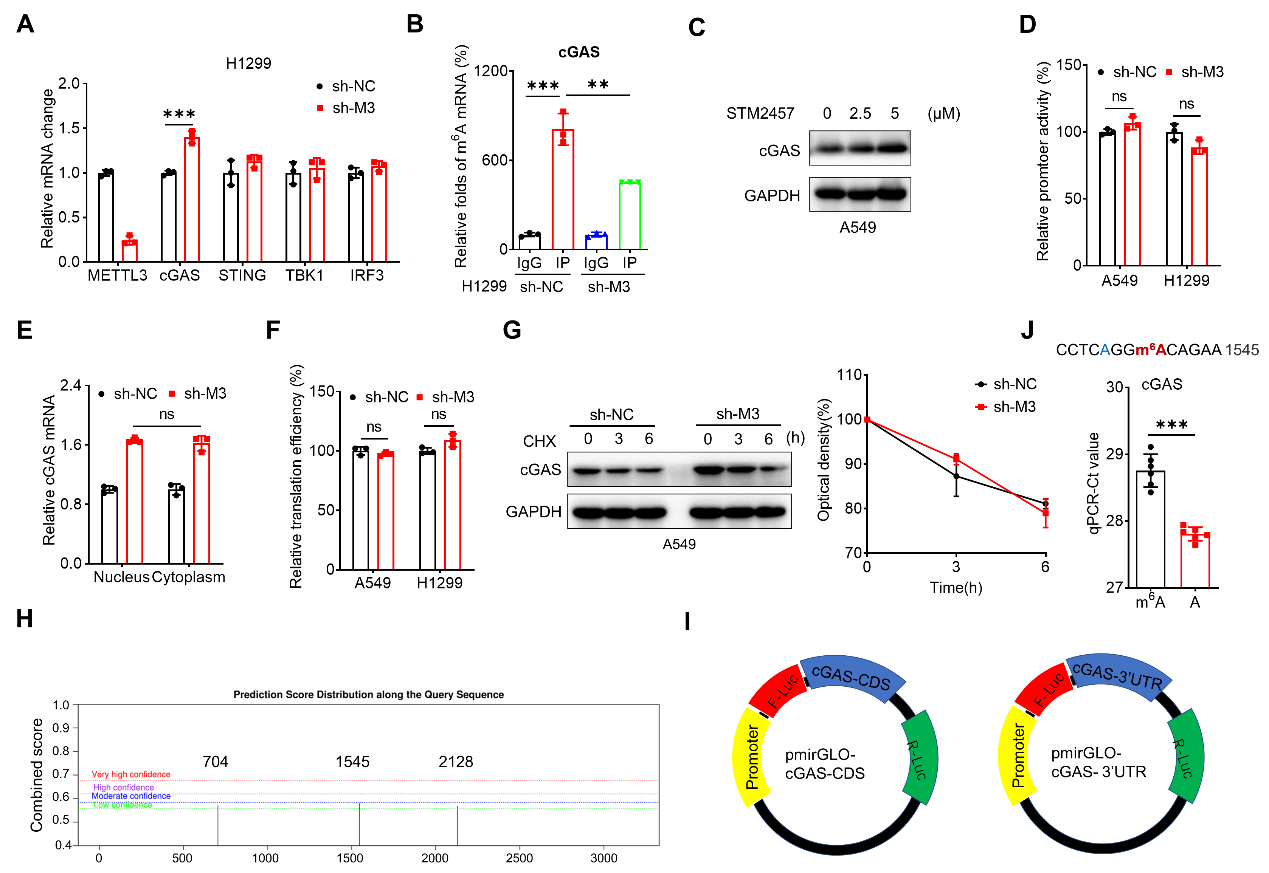

**Supplementary Figure 5 METTL3 destabilized cGAS mRNA via binding YTHDF2 and A1545 methylation**

(A) The mRNA expression of cGAS, STING, TBK1, IRF3 in sh-control and sh-METTL3 H1299 cells;
(B) m^6^A RIP-qPCR analysis of cGAS in sh-control and sh-*METTL3* H1299 cells;
(C) The protein expression of cGAS in A549 cells treated with STM2457;
(D) Cells were transfected with pGL3-Basic- cGAS -luc reporter and pRL-TK plasmid for 24 h, the promoter activities were presented as the ratios of the reporter normalized to pRL-TK plasmid;
(E) The relative levels of nuclear versus cytoplasmic cGAS mRNA in sh-control and sh-*METTL3* A549 cells;
(F) The translation efficiency of endogenous cGAS in sh-control and sh-*METTL3* A549 cells;
(G) Cells were treated with 10 μg/ml CHX for the indicated time periods, the protein expression of cGAS was detected by western blot analysis (left) and quantitatively analyzed (right);
(H) The predicted m^6^A peaks in *MSH5* mRNA from the m^6^A sites predictor SRAMP;
(I) Schematic representation of pmirGLO-*cGAS* CDS and pmirGLO-*cGAS* 3’UTR reporter;
(J) The threshold cycle (Ct) of qPCR showing SELECT results for detecting m^6^A site in the potential m^6^A site (A1545) and negative A site (A1542) of cGAS in A549 cells;

Data are presented as mean ± SD from three independent experiments. **p*<0.05, ***p*<0.01, ****p*<0.001, ns, no significant, by Student’s *t* test between two groups and by one-way ANOVA followed by Bonferroni test for multiple comparison.

**Materials and methods**

1. **Cell line and cell culture**

Human A549, H1299 and murine LLC cells were commercially obtained from American Tissue Cell Culture (ATCC, USA) and maintained by our laboratory. Cells were cultured in high-glucose DMEM (GIBIO, USA) medium supplemented with 10% FBS (GIBIO, USA) and 100 U/ml penicillin/streptomycin (Beyotime, China) under an atmosphere of 5% CO2 at 37℃. The stable cell lines were cultured in medium containing puromycin or neomycin until 3 days before experiment.

2. **Plasmid, siRNA, shRNA and generation of stable cell lines**

The cDNA of MSH5 were cloned into the pcDNA3 vector (Invitrogen, USA), and the CDS of METTL3 was cloned into the ppB vector to generate over expression plasmid, while the METTL3 mutant DA (D395A) plasmid was generated in our previous study^[1]^. The following siRNAs were synthesized (Ribobio, China) and used in the study: siRNA negative control (si-NC): 5’-UUC UCC GAA CGU GUC ACG U-3’; IGF2BP2 #1: 5’-CAT GCC GCA TGA TTC TTG A-3’; IGF2BP2 #2: 5’-GAA CGA ACT GCA GAA CTT A-3’; IGF2BP2 #3: 5’-AAC AGG GAC CAA GAT AAC A-3’; YTHDF2 #1: 5’-GAC CAA GAA TGG CAT TGC A-3’; YTHDF2 #2: 5’- GCA CAG AAG TTG CAA GCA A-3’. To generate stable cell lines with continuous suppression of METTL3, cells were transfected with lentivirus-shRNA for negative control, and METTL3, respectively, before selection with puromycin.

3. **Human tissue microarray and immunohistochemical (IHC) analysis**

The study was approved by the Institutional Ethics Committee of Sun Yat-sen University (No. YBL2024157). Paired tumor and adjacent non-tumor paraffin tissue microarray for human lung carcinoma were purchased from YEPCOME Biotechnology Co., Ltd (Shanghai, China). The microarray comprised of 50 pairs of LUAD and corresponding non-tumor paraffin tissue samples. IHC analysis was performed as previously described.

4. **Flow cytometric analysis**

Cell surface marker staining and flow cytometric analysis for CD3, CD4, CD8, CD45 (Biolegend) expression was performed as described previously^[2]^. Cy5.5-anti-Mouse CD3e (65-0031-U025, TONBO Bioscience, USA), APC-anti-Mouse CD4 (25-0042-U025, TONBO Bioscience, USA), cy7-anti-Mouse CD8a (60-0081-U025, TONBO Bioscience, USA), FITC- anti-Mouse CD45 (35-0451-U025, TONBO Bioscience, USA).

5. **Western blot analysis**

Cells were washed with PBS, lysed in radio-immunoprecipitation assay (RIPA) buffer containing 1 mM PMSF (Beyotime, China), and placed on ice for 30 min. Then, cells were centrifuged at 12,000×g for 20 min, and the protein concentration was determined using BCA Protein Assay Kit (Thermo Fisher, USA). Proteins were separated by 10% SDS-PAGE gel and electro-transferred to polyvinylidene difluoride membrane (Bio-Rad, USA). The membranes were blocked with 5% nonfat milk in 1×PBST for 30 min at room temperature, and incubated at 4℃ overnight with the following primary antibodies: anti-METTL3 (15073-1-AP, proteintech, China); anti-YTHDF2 (ab220163, Abcam, England); anti-YTHDF1 (ab99080, Abcam, England); Anti-IGF2BP1 (8482S, CST, USA); Anti-IGF2BP2 (14672S, CST, USA); Anti-IGF2BP3 (25864S, CST, USA); Anti-γH2AX (2577, CST, USA); Anti-cGAS (A25686, Abclonal Technology, China); Anti-STING (A21051, Abclonal Technology, China); Anti-p-STING (AP1369, Abclonal Technology, China). Anti-GAPDH (5174Ss, CST, USA) was used as an internal loading control. After incubation with corresponding secondary antibodies (CST, USA), the membranes were incubated with ECL substrate (Thermo Fisher, USA).

6. **RNA-extraction and real-time PCR**

Total RNA was isolated using TRIzol Reagent (Agbio, AG21102, China) and reversed by Evo M-MLV (Agbio, AG11706, China). qRT-PCR was assessed with SYBR Green II (Agbio, AG11701, China) using CFX Manager 3.1 (Bio-Rad, USA) as recommended by the manufacturer’s protocol. The primers used are listed at Table S5 in the Supplementary Data. GAPDH was used as a control for normalization. The relative gene expression levels were calculated using 2^−ΔΔCT^ method.

7. **m^6^A-RIP qPCR**

We performed m^6^A qPCR using Magna MeRIP™ m^6^A Kit (Millipore, MA) in accordance with manufacturer’s protocol. Briefly, 200 μg total RNA was isolated and randomly fragmented with chemical reagents treatment followed by the immunoprecipitation with 5 μg m^6^A antibody or mouse IgG which was linked to Magna ChIP Protein A/G Magnetic Beads. After extensive washing with IP Buffer, the beads were treated with proteinase K for 30 min at 55℃ with occasional shaking. RNA was purified from the supernatant using TRIzol Reagent following the manufacturer’s instructions. Interested mRNA levels in the elutes were measured by RT-qPCR.

8. **RIP-RT-PCR**

Two 10-cm plates of cells were washed twice with cold PBS before collected. 400 μl IP lysis buffer (150 mM KCl, 25 mM Tris (pH 7.4), 5 mM EDTA, 0.5 mM DTT, 0.5% NP40, 1×protease inhibitor, 1 U/μl RNase inhibitor) was added and resuspended it on ice. The lysate was centrifuged at 12,000×g for 10 min. Then Magnetic beads pre-coated with 4 μl targeted antibodies or mouse IgG (NEB, USA) were incubated with sufficient cell lysates at 4℃ overnight. The beads containing immunoprecipitated RNA-protein complex were treated with proteinase K to remove proteins. Then interested RNAs were purified by TRIzol methods and detected by RT-qPCR with the normalization to input.

9. **Sub-cellular fraction**

Fractionation of nuclear and cytoplasmic samples was conducted using Nuclear and Cytoplasmic Extraction Kit (Beyotime, China) according to the manufacturer’s guidelines.

10. **Protein stability**

To measure protein stability, cells were seeded in 6-well plates and treated with cycloheximide (CHX, Catalog #14126, Cayman, USA) at final concentration 20 μg/ml during indicated times. Cells were collected and lysed in lysis buffer. The expression of proteins was measured through western blot analysis.

11. **mRNA stability**

To measure RNA stability in tumor cells, actinomycin D (Act-D, Catalog #A9415, Sigma, USA) at 10 μg/ml was added to cells in 6-well plates. After incubation at the indicated times, cells were collected, and RNA was isolated for real-time PCR. Half-life (t_1/2_) of mRNA were calculated using ln2/slope and 18S was used for normalization.

12. **SELECT**

SELECT qPCR was conducted by following Xiao’s protocol ^[3]^ and our previous study^[4]^. Briefly, 1500 ng of total RNA was mixed with 40 nM up and down primers and 5 μM dNTP in 17 μl 1×CutSmart buffer (NEB, China). The mixture was incubated with the follow program: 90℃ for 1 min, 80℃ for 1 min, 70℃ for 1 min, 60℃ for 1 min, 50℃ for 1 min and 40℃ for 6 min. The sample was further mixed with 0.5 U SplintR ligase, 10 nM ATP and 3 μl of 0.01 U Bst 2.0 DNA polymerase and incubated at 40℃ for 20 min and denatured at 80℃ for 20 min. Afterwards, 20 μl qPCR reaction containing 2 μl of final reaction mixture, 2×SYBR Green Master Mix (Takara, Japan), and 200 nM SELECT primers was performed. The qPCR program was 95℃, 5 min; (95℃, 10 s; 60℃, 35 s)×40 cycles; 95℃, 15 s; 60℃, 1 min; 95℃, 15 s; 4℃ hold. Results were calculated by normalized the Ct values of samples to their corresponding Ct values of control. All assays were performed with three independent experiments.

13. **Database (DB) search**

We used the TIMER2.0 web server (http://timer.comp-genomics.org/) to investigate the correlation between METTL3 expression and immune cell infiltration. Correlations between METTL3 protein expression in LUAD tissues and other proteins such as cGAS were extracted from CPTAC database. The mRNA expression of METTL3 and MSH5 in LUAD and normal tissues were analyzed in the TCGA database.
